# Enantiomers of Chloroquine and Hydroxychloroquine Exhibit Different Activities Against SARS-CoV-2 *in vitro*, Evidencing *S*-Hydroxychloroquine as a Potentially Superior Drug for COVID-19

**DOI:** 10.1101/2020.05.26.114033

**Authors:** Guanguan Li, Jing Sun, Yi-You Huang, Yingjun Li, Yongjie Shi, Zhe Li, Xiang Li, Feng Hua Yang, Jincun Zhao, Hai-Bin Luo, Tony Y. Zhang, Xumu Zhang

## Abstract

In all of the clinical trials for COVID-19 conducted thus far and among those ongoing involving chloroquine or hydroxychloroquine, the drug substance used has invariably been chloroquine (CQ) diphosphate or hydroxychloroquine (HCQ) sulfate, i.e., the phosphoric or sulfuric acid salt of a racemic mixture of *R*- and *S*-enantiomer (50/50), respectively. As a result, the clinical outcome from previous CQ or HCQ trials were, in fact, the collective manifestation of both *R* and *S-* enantiomers with inherent different pharmacodynamic and pharmacokinetic properties, and toxicity liabilities. Our data for the first time demonstrated the stereoselective difference of CQ and HCQ against live SARS-CoV-2 virus in a Biosafety Level 3 laboratory. *S*-chloroquine (*S*-CQ) and *S*-hydroxychloroquine (*S*-HCQ) significantly more active against SARS-CoV-2, as compared to *R*-CQ and *R*-HCQ, respectively. In addition, M^pro^, as one of the critical enzymes for viral transcription and replication, also exhibited an enantioselective binding affinity toward the *S*-enantiomers. The most significant finding from this study is the pronounced difference of the two enantiomers of CQ and HCQ observed in hERG inhibition assay. The IC_50_ value of *S*-HCQ was higher than 20 μM against hERG channel, which was much less active over all tested CQ and HCQ compounds. Moreover, *S*-HCQ alone did not prolong QT interval in guinea pigs after 3 days and 6 days of administration, indicating a much lower cardiac toxicity potential. With these and previous findings on the enantio-differentiated metabolism, we recommend that future clinical studies should employ *S*-HCQ, substantially free of the *R*-enantiomer, to potentially improve the therapeutic index for the treatment of COVID-19 over the racemic CQ and HCQ.

## INTRODUCTION

In the last few months, the virulence and lethality of coronavirus breakout have presented an unprecedented challenge to the medical community and world governments. The coronavirus disease 2019 (COVID-19), caused by severe acute respiratory syndrome coronavirus 2 (SARS-CoV-2) ^1–2^ has become a global pandemic. It has now affected more than 188 countries and regions, ^1, 3–6^ becoming a global pandemic as declared by the World Health Organization (WHO). By August 21^st^, 2020, the cumulative number of confirmed cases of COVID-19 infection has exceeded 22.6 million worldwide, with more than 793,710 recorded deaths (a fatality rate of 3.5 % approximately), according to real-time data released by Johns Hopkins University.^7^

With the prospect of effective vaccines months away from being developed and launched, scientists and physicians alike have turned to existing drugs with known toxicity and human pharmacokinetic data to deal with this global healthcare crisis. Numerous treatment regimens for COVID-19 have been proposed and tested in clinical trials with varying degrees of controls and sizes of subject enrollment. With limited access to compounds and live virus for testing, research institutions around the world have also resorted to tools like virtual screening and modeling to compile lists of potential drugs that may be efficacious for COVID-19.^8–10^ Many of these have been evaluated against SARS-CoV-2 *in vitro*. These include some medications commonly used for decades, such as chloroquine,^11–13^ arbidol,^14^ ribavirin,^15–17^ lopinavir, etc.,^18–20^ The list also contains some drugs in development, for instance, remdesivir,^11, 17^ a previously studied drug candidate for treating Ebola infection.^21–22^

Among the most notable and also controversial repurposed drugs are chloroquine (CQ) and hydroxychloroquine (HCQ) as shown in Figure 1A, with several initial promising results especially when combined with azithromycin or zinc supplement being reported,^23–31^ only to be followed by contradicting reports^32–37^ on lack of efficacy and presence of severe side effects, especially among the higher dosed patients. Based on the preclinical results, both CQ and HCQ have displayed inhibition of viral replication.^11–13^ The 70-year-old CQ was first used in clinical practice as an anti-malarial drug, and its indications were extended to treat systemic lupus erythematosus, rheumatoid arthritis, and several infections.^38^ Originally developed as a safer alternative to CQ for treating malaria, the analogous compound hydroxychloroquine sulfate^39^ has become a popular drug for treating autoimmune diseases including rheumatoid arthritis (RA) and system lupus emphysema (SLE).^40–41^ In fact, it is the 2^nd^ most prescribed medicine for rheumatoid arthritis in the United States, with nearly half a billion tablets dispensed in 2019. Both drugs have a long history as inexpensive and well-received treatment options for malaria, autoimmune, as well as infectious diseases, even for women patients during pregnancy.^38^ We believe the verdict is still out on their ultimate safety and efficacy. By the last account, there are currently 41 and 133 ongoing or planned clinical trials (clinicaltrials.gov) against COVID-19 involving the use of CQ and HCQ, respectively.

**Figure 1.**
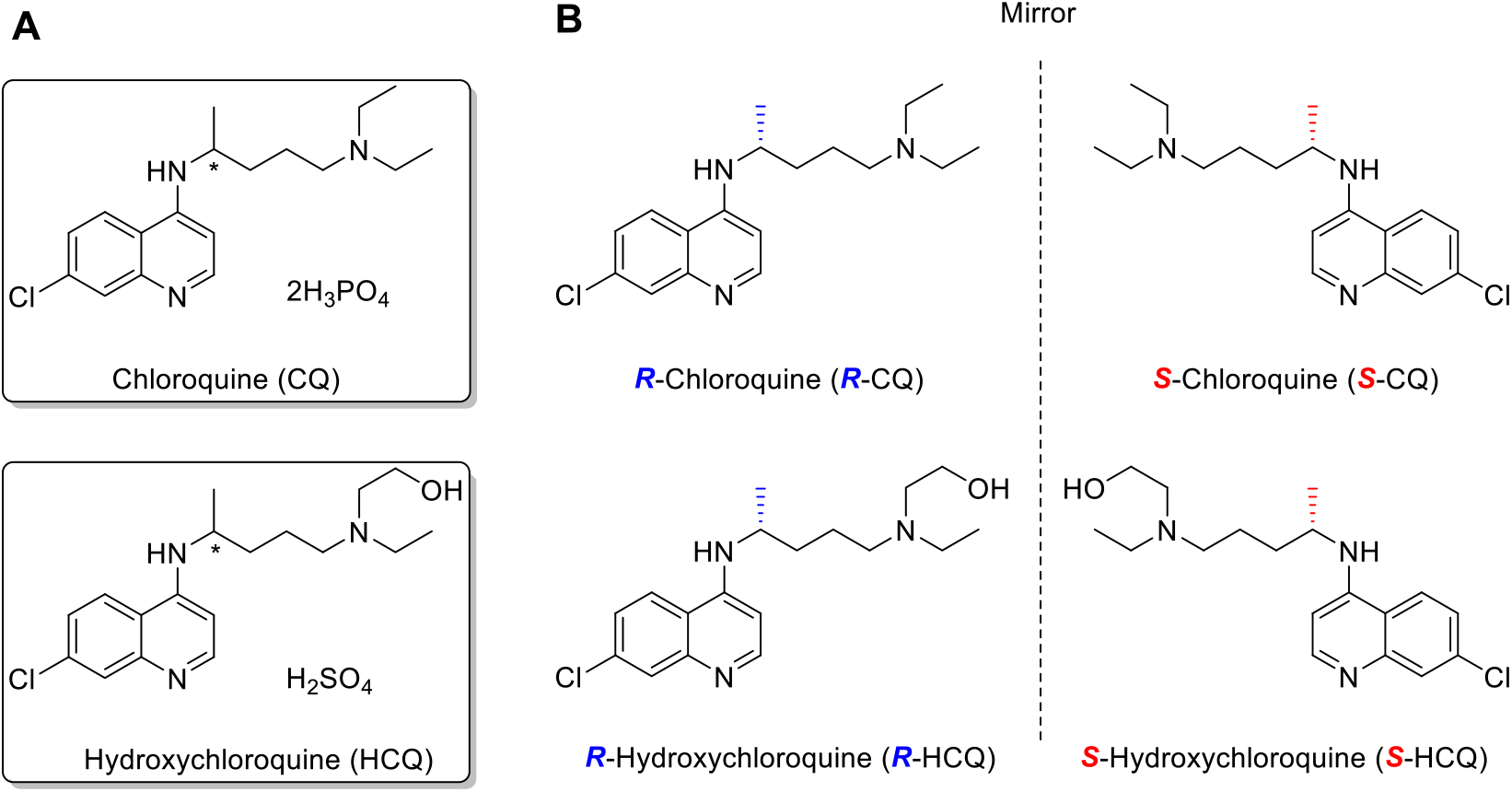
Structures of chloroquine and hydroxychloroquine (A), as well as their enantiomers as mirror images to each other (B).

While remdesivir, another repurposed antiviral that has also generated enormous excitement in the global communities, has also been approved by FDA under an EUA for treating COVID-19 and is emerging as the standard of care, its demonstrated efficacy is far from ideal. The development of less expensive, safer, more effective, and more accessible treatment options is still an urgent unmet medical need.

We noticed certain caveats about CQ and HCQ that are worthy of being taken into accounts by stakeholders. First, CQ and HCQ have been in use for decades by millions of patients with diverse health conditions since their approval in the mid-1900s; hence, a wealth of information about the adverse events is available.^42–44^ Secondly, unlike i.v. administered drugs including remdesivir, CQ and HCQ can be taken conveniently as oral tablets. Thirdly, CQ and HCQ are among the least expensive drugs being considered for COVID-19, with an excellent prospect of accessibility for the global population should its safety and efficacy be proven by further and more definitive, randomized, and controlled clinical trials with more appropriate tailoring according to patient conditions, time of intervention, and dosing regimen. Last but not least, all preclinical and clinical trials of CQ and HCQ for COVID-19 thus far have been carried out using the common CQ and HCQ formulations,^45^ *i.e*., the phosphoric or sulfuric acid salt of a ***racemic*** mixture of CQ or HCQ, respectively.

Commercial chloroquine and hydroxychloroquine drug products are administrated as their respective racemates of a 50:50 mixture of two enantiomers: *S*- and *R*-isomers. The *R*- and *S*- isomers are in an enantiomeric relationship, meaning that these two compounds have the same chemical compositions but are a mirror image relationship to each in the spatial arrangement, as shown in Figure 1B. Compounds that cannot superimpose with their own mirror images are defined as chiral compounds. Life, the biological system, is chiral in nature, as proteins are composed of amino acids of predominately that of the *L*-configuration, and sugars of mostly *D*-configurations. This predisposes different enantiomers of a chiral drug to interact with proteins differently, and hence the different pharmacological effects would be expected. As chiral drugs, the two enantiomers of CQ or HCQ, might exhibit distinct efficacy and safety profiles in patients. These differences, big or small, can translate into performance variances that could potentially influence the decision process on whether either enantiomer of CQ or HCQ can ultimately become a viable therapeutic agent for COVID-19 through further testing, including clinical trials. The clinical trials for COVID-19 conducted so far have all been using a racemic mixture of either CQ or HCQ; hence the observations were actually the collective manifestation of the two different albeit very similar optical isomers of these drugs. CQ, as racemic mixtures, exhibited some antiviral activity against SARS-CoV-2 *in vitro* (EC_50_= 1.13 μM in Vero E6 cells); HCQ was also active with less toxicity;^11-13, 45^ many clinical reports also presented that CQ and HCQ prevented patients from experiencing severe symptoms;^24–31^ however, the efficacy and possible toxicity of each individual isomer against SARS-CoV-2 have not been reported in any preclinical or clinical studies yet. In this paper, we wish to present the preliminary results on such a difference exhibited by the enantiomers of CQ and HCQ, respectively, when subjected to *in vitro* test against SARS-CoV-2.

## RESULTS

### Chiral HPLC separation of both chloroquine and hydroxychloroquine

From the chiral HPLC separation of the racemic chloroquine, the first eluting compound at 4.86 minutes was found to be dextrorotary (+) and thus assigned as the *S*-CQ and the second compound at 5.33 minutes as *R*-CQ, with a specific rotation of 79.7 and −74.4, respectively, as depicted in Figure 2D-F. For the case of hydroxychloroquine in Figure 2A-C, the first eluting compound at 10.17 minutes was determined as *S*-HCQ, and the second compound at 11.85 minutes was *R*-HCQ, with a specific rotation of 95.6 and −107.8, respectively. The enantiomeric excess value of each enantiomer for both HCQ and CQ was found to be higher than 98%, as shown in Figure 2B, C, E, and F. In all cases, baseline separation was achieved. Material recovery was 8.32 g for loaded 9.64 g of HCQ case, and 5.04 g for loaded 5.04 g of CQ.

**Figure 2.**
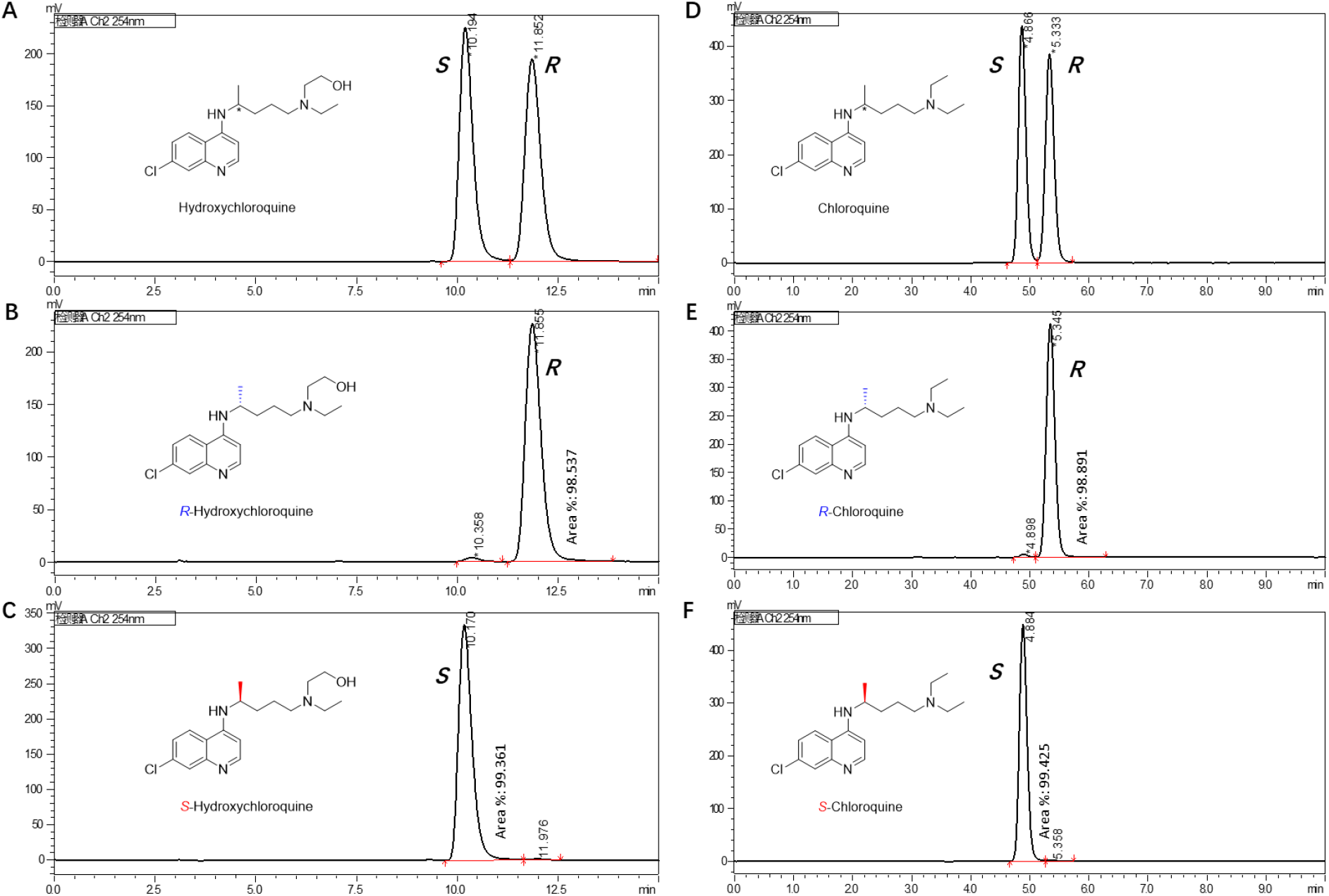
Chromatograms for the chiral HPLC separation from the racemic hydroxychloroquine (A-C) and chloroquine (D-F). Optimal chromatographic conditions for the two racemic mixtures: mobile phase, isocratic *n*-hexane/*iso*propanol/diethylamine with a ratio of 85:15:0.1 (v/v/v); eluent flow rate, 1.0 mL/min; column temperature, 35 °C; detection wavelength, 254 nm.

### Antiviral activity in vitro

Results from the *in vitro* antiviral activity against SARS-CoV-2 in Vero E6 cells showed that both CQ and HCQ and their enantiomers exhibited respectable antiviral effect in a concentration-dependent manner. The IC_50_ values for the racemic chloroquine diphosphate (Rac-CQ) and its *R* and *S* enantiomer against SARS-CoV-2 *in vitro* were 1.801 μM, 1.975 μM, and 1.761 μM, respectively, as displayed in Figure 3A. Shown in Figure 3B, the IC_50_ values for the racemic hydroxychloroquine sulfate (Rac-HCQ) and its *R* and *S* enantiomer were 1.752 μM, 2.445 μM, and 1.444 μM, respectively. In addition, we tested the antiviral activity of azithromycin that was used as a combination drug with HCQ for treating COVID-19 in patients.^46^ The single-dose of azithromycin afforded an IC_50_ value of 14.57 μM, shown in Figure 3C. The results from immunofluorescence microscopy of SARS-CoV-2 infection displayed a better antiviral effect of the *S*-enantiomers of both CQ and HCQ, as compared to their *R*-enantiomer starting at the dose of 1 μM, respectively. The data showed that the antiviral activity of *S*-CQ was were relatively similar to the racemic CQ (Figure 4A), whereas the efficacy of the *S*-HCQ was much higher than its racemates (Figure 4B). On the other hand, azithromycin alone started to exhibit an antiviral effect at a relatively high concentration, which is over 10 μM (Figure 4C).

**Figure 3.**
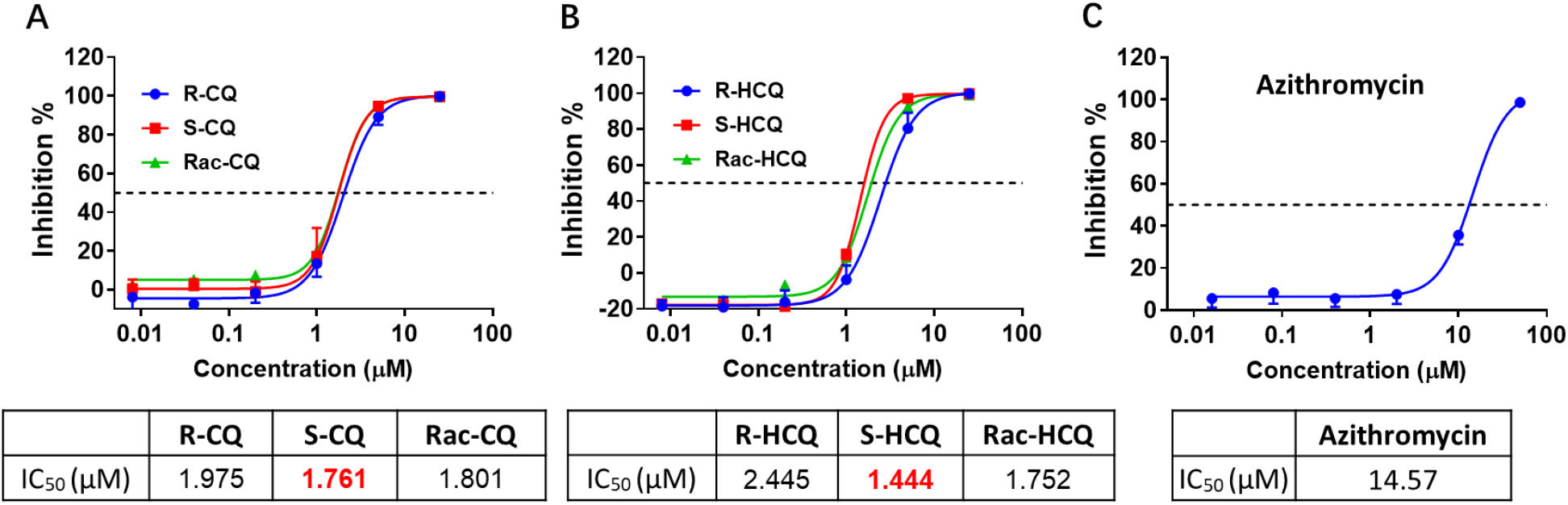
The antiviral activities of racemic and enantiomeric chloroquine diphosphate (A) and hydroxychloroquine sulfate (B), as well as azithromycin (C) against SARS-CoV-2 *in vitro*. Vero E6 cells were infected with SARS-CoV-2 (MOI = 0.05) at different concentrations: 0.008, 0.04, 0.2, 1, 5, and 25 μM for CQ and HCQ for 24 h (concentrations at 0.008, 0.04, 2, 10 and 50 μM for AZM). Data represented are the mean value of % inhibition of SARS-CoV-2 on Vero E6 cells.

**Figure 4.**
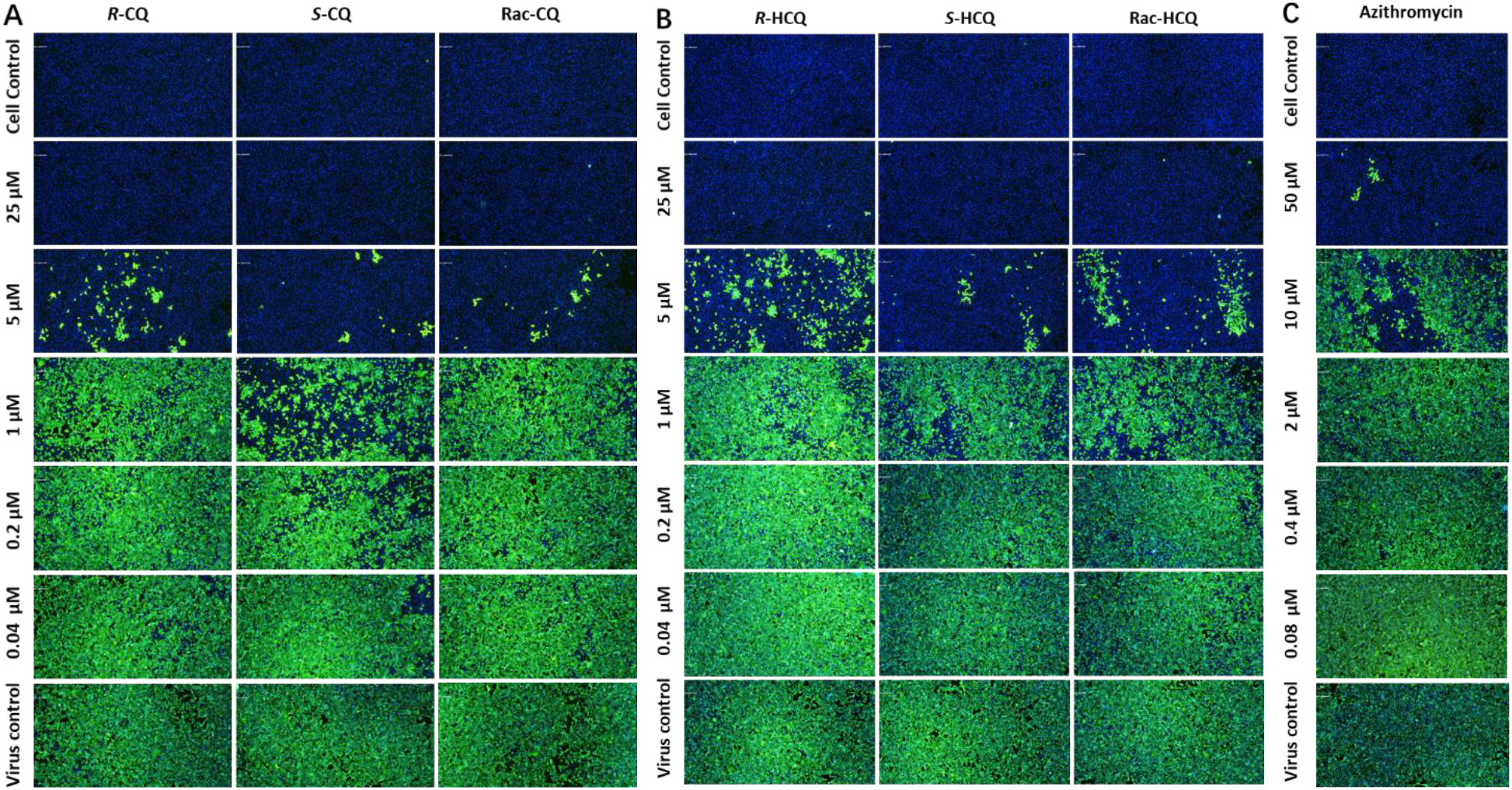
Immunofluorescence microscopy of virus infection with the treatment of chloroquine diphosphate (A) and hydroxychloroquine sulfate (B), as well as their enantiomers, and azithromycin (C) against SARS-CoV-2 *in vitro*. Experiments of virus infection and drug administration were conducted, as described in Figure 3. At 24 h p.i., the infected cells were fixed and then incubated with the primary antibody (rabbit polyclonal antibody against SARS Coronavirus Nucleoprotein), followed by Alexa 488 labeled secondary antibody. The nuclei were stained with DAPI. Bars, at 100 μm.

The most significant finding is that both *S*-CQ and *S*-HCQ exhibited more pronounced activities (+31 % and + 60 %, respectively, based on the mean value n = 6, see Figure S13 and S14 for additional data) than their respective *R*-enantiomer against SARS-CoV-2 *in vitro*. The IC_50_ value of the racemic mixtures of CQ and HCQ fell between the two enantiomers, as expected, which indicated that, at the testing conditions, the *S*-isomers were the major contributors to the antiviral activities of the racemates. It implies that the *R*-isomers were either coming along for the ride, being minor contributors to the antiviral activities, providing hitherto unknown benefits such as cytokine storm management, or contributing to untoward side effects.

### hERG inhibition

The most concerning side effects associated with CQ and HCQ applications relate to cardiovascular toxicity, as confirmed by recent trials for COVID-19. These drugs are known to increases the QT interval^47^ by blocking the hERG (human ether-a-go-go related gene, Kv11.1) channel, one of the major ion channels conducting potassium ions out of the heart muscle cells. Limited data indicated that *S*-CQ seemed to be less toxic than the *R*-isomer.^43^ Toward elucidating each enantiomer’s individual contribution to cardiovascular liabilities, we subjected them toward hREG channel assay, as an *in vitro* predictive test for cardiovascular side effect surrogate for *in vivo* test, ^48^ with results shown in Table 1.

**Table 1.** The IC_50_ values of CQ, HCQ, and their enantiomers against hERG ion channel.

| Compound | Maximum concentration inhibition (%) | Maximum concentration (μM) | IC<sub>50</sub> (μM) |
| --- | --- | --- | --- |
| Cisapride* | 99.5 | 3 | 0.041 |
| Rac-CQ | 86.1 | 40 | 4.56 |
| *R*-CQ | 89.8 | 40 | 4.83 |
| *S*-CQ | 72.3 | 40 | 12.8 |
| Rac-HCQ | 72.0 | 40 | 12.8 |
| *R*-HCQ | 70.9 | 40 | 15.7 |
| ***S*-HCQ** | **32.8** | **20** | **>20** |

\*Cisapride was used as a positive control for hERG ion channel assay.

The hERG data predicts for a superior cardiovascular safety profile for *S-*HCQ and *S-*CQ over their respective *R-*isomers. As shown in Table 1, the IC_50_ values of racemic CQ and HCQ were 4.56 and 12.8 μM, respectively. The IC_50_ values of *R-*CQ and *S-*CQ were 4.83 and 12.8 μM. Similarly, the hERG activity for *R-*HCQ and *S-*HCQ were 15.7 and > 20 μM, respectively. The *S-*isomers of both CQ and HCQ were uniformly less active than their *R-*isomers at inhibiting hERG. Though an hERG IC_50_ over 10 μM may be considered acceptable for a drug in early development, the extremely long half-life and the large volume of distribution of HCQ must be taken into account. The lower activity of *S-*HCQ at inhibiting hERG (IC_50_ >20 μM) suggests that administering an enantiomerically pure *S*-HCQ presents less a chance for cardiac toxicity, as compared with racemic CQ and HCQ.

### QT assessment in vivo

Although *S*-HCQ exhibited an acceptable IC_50_ value in the *in vitro* hERG binding affinity assay, we elected to conduct a more definitive cardiovascular toxicity test, *in vivo* animal QT prolongation for this enantiomer, as International Conference on Harmonization (ICH) Guidelines stipulates that the drug candidates are required to pass the *in vivo* QT assessment for cardiac safety evaluation. Thus, we used an anesthetized guinea-pig assay ^49^ for a practical and reliable prediction of QT interval prolongation assay, since the cardiac action potential and electrocardiogram of these small animals are similar to those of humans. The treatment dosage for guinea pig was adjusted and converted by body weight according to the clinical dosage of humans ^50^. As displayed in Figure 5 and Table 2, the results showed that there was no significant difference in △QTc ^51^ between the *S*-HCQ group and the control group after 3 days and 6 days of administration (p = 0.4146 and p = 0.3401, respectively). However, when combined with AZM, the change of △QTc was significantly increased (p = 0.0261 and p = 0.0114, respectively) in both day 3 and day 6, indicating that the prolongation of QTc was related to AZM. Moreover, there were significant differences between *S*-HCQ group and *S*-HCQ plus AZM group (p<0.001 and p<0.001, respectively).

**Figure 5.**
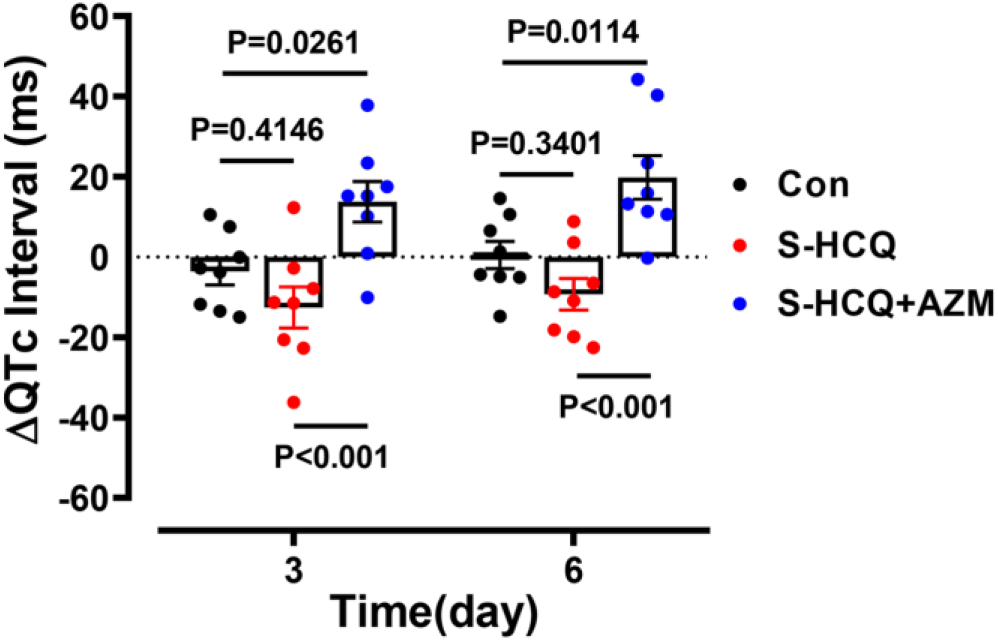
Differences of QTc *in vivo* at before or after 3 days and 6 days using *S*-HCQ with or without AZM. △QTc, change in the corrected QT interval. Data are presented as the mean ± S.E.M.

**Table 2.** Quantitative assessment of the effect of *S*-HCQ with or without AZM on the QTc prolongation *in vivo*.

| Time (day) | ΔQTc (ms) | | |
| --- | --- | --- | --- |
|  | Control | *S*-HCQ | *S*-HCQ + AZM |
| 3 | $-3.625 \pm 3.355$ | $-12.59 \pm 5.113$ | $13.75 \pm 5.066$ |
| 6 | $0.475 \pm 3.423$ | $-9.3 \pm 3.949$ | $19.83 \pm 5.425$ |

Data are presented as the mean ± S.E.M (n= 8 for each group). $\Delta$ QTc represents the difference of QTc before and after administration.

### SARS-CoV-2 main protease (M^pro^) inhibition

We first screened CQ, HCQ and their enantiomers and found that they indeed bound to M^pro^ with various degrees of affinity. Perturbation calculations predicted binding free energies (ΔG_pred_)^52^ of M^pro^ with *R*- and *S-*HCQ to be −9.8 and −11.3 kcal/mol, respectively in Table 3. The computationally modeled structure of *S-*HCQ binding with SARS-CoV-2 M^pro^ is depicted in Figure 6A, which shows the roles of key residues of the protease for binding with *S-*HCQ, including His 41, Val 42, Cys 44, Met 49, Met 165, and Asp187, Gln 189, and Thr190.

**Table 3.** Enzymatic assay of CQ, HCQ, and their enantiomers targeting M^pro^.

| Compound | IC<sub>50</sub> (μM) | Converted<br>K<sub>i</sub> (μM) | ΔG<sub>exp</sub><br>(kcal/mol) | ΔG<sub>pred</sub><br>(kcal/mol) |
| --- | --- | --- | --- | --- |
| *R*-CQ | 8.36 ± 0.39 | 1.03 ± 0.05 | -8.2 | -10.0 |
| *S*-CQ | 5.27 ± 0.49 | 0.65 ± 0.06 | -8.4 | -10.6 |
| Rac-CQ | 7.54 ± 0.25 | 0.93 ± 0.03 | — | — |
| *R*-HCQ | 3.26 ± 0.38 | 0.40 ± 0.05 | -8.7 | -9.8 |
| ***S*-HCQ** | **2.47 ± 0.39** | **0.30 ± 0.05** | **-8.8** | **-11.3** |
| Rac-HCQ | 2.77 ± 0.41 | 0.34 ± 0.05 | — | — |

K<sub>i</sub> = IC<sub>50</sub>/(1+[S]/K<sub>m</sub>), [S] = 10 μM, K<sub>m</sub> = 1.4 μM. ΔG<sub>exp</sub> = RT ln (K<sub>i</sub>)

**Figure 6.**
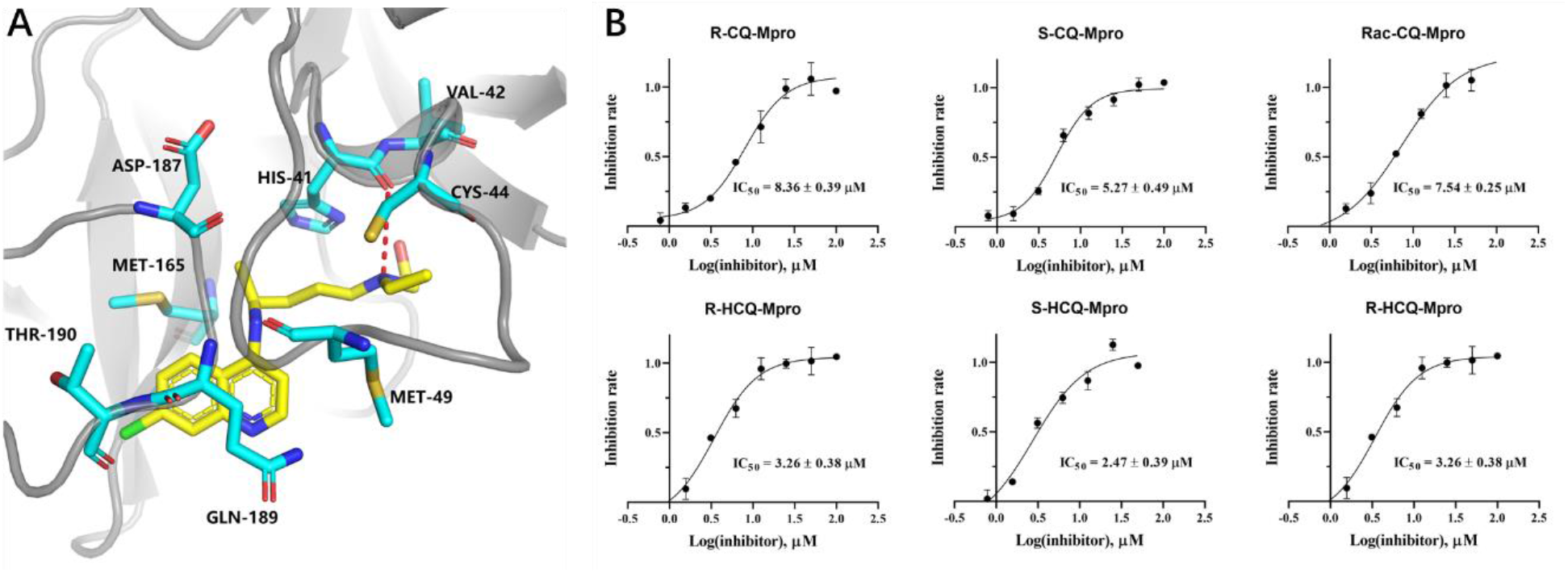
**A**.The putative binding pattern between *S-*HCQ and SARS-CoV-2 M^pro^ after MD simulation. *S-*HCQ is shown in yellow in the stick model, the key amino acid residues of M^pro^ are shown in cyan in the stick model, and the hydrogen bond is shown as red dashed line. **B**. Inhibitory curves of enantiomeric or racemic chloroquine diphosphate and hydroxychloroquine sulfate against SARS-CoV-2 M^pro^. IC_50_ values and error bars are represented by mean ± SD based on three independent measurements.

We then subjected the enantiomers of CQ and HCQ to M^pro^ enzymatic inhibition studies. Indeed, *S-*HCQ appeared as the most potent isomer with a Ki of 0.30 μM, as compared with that of 0.40 μM of *R-*HCQ (Figure 6B). The same trend was observed for *S-*CQ and *R-*CQ, though their Ki values were higher than the HCQ series. Among all 6 compounds tested, summarized in Table 3, *S-*HCQ showed the highest potency with an IC_50_ value of 2.47 μM.

## DISCUSSION

### Chiral drugs and performance difference of enantiomers

*R*- and *S*- enantiomers are different substances and by definition should not be expected to have the same biological effect in a living system, which is inherently chiral. Enantiomers often have different pharmacodynamics and pharmacokinetics behaviors and hence different therapeutic efficacy and safety profiles. However, the significance of this principle in drug development has not been fully appreciated at the time when CQ and HCQ were developed 70 some years ago.

One of the most notorious examples of drug chirality is thalidomide, a chiral drug first marketed as a racemic form in 1957 in Europe as an antiemetic for treating morning sickness and other ailments. The drug was withdrawn from the market in 1961 amid the emergence of birth defects among newborn children of woman patients. It was later determined that the two enantiomers caused distinctly different effects from one another, and the *S*-isomer was attributed to being the culprit for the teratogenic properties.^53^ Since the enantiomers of thalidomide interconverts *in vivo*, efforts to develop a stable, single-enantiomer thalidomide was thwarted. Eventually, the racemic form was repurposed and launched for oncology applications under the brand Thalomid, strictly contraindicated for pregnancy. The thalidomide tragedy brought up a lot of attention and concerns about the importance of chirality in the drug development process. This eventually culminated in more specific guidance and regulations regarding chiral drug development by the FDA in the 1990s. CQ or HCQ would unlikely be approved as they had been submitted to a current regulatory authority, who will likely demand the delineation of each enantiomer’s properties and demonstration of the non-inferiority of the racemate in the context of safety and efficacy before it is allowed for marketing approval.

Unlike thalidomide, CQ and HCQ are stereochemically stable and the interconversion between the two enantiomers has not been observed *in vivo* and is unlikely from a mechanistic point of view. The separation of the *R*- and *S-*enantiomers of CQ and HCQ has not been reported until the early 1990s,^54–55^ as part of the metabolite profiling studies for HCQ.^54^ With the enantio-discriminating analytical methods available, for CQ, HCQ and their respective metabolites, several studies demonstrated varying degrees of differences of pharmacological properties between the two enantiomers of CQ and HCQ.

### Pharmacokinetics, pharmacology, and toxicology aspect of CQ and HCQ enantiomers

Both CQ and HCQ have extremely long half-lives approximating 50 days, predominantly due to their huge volume of distribution.^56^ With such a low metabolic rate, the drugs accumulate excessively in tissues over time, especially eyes, skin, brain, heart, and liver. Long term and higher doses other than the recommended maximal dose for chloroquine and hydroxychloroquine (300 mg/kg and 400 mg/kg, respectively, according to the FDA) put the patients at risk of adverse effects such as irreversible retinopathy, neuropathic, and cardiac toxicities.^38, 40–41, 57^ One of the major adverse effects associated with the clinical applications of CQ and HCQ is ocular toxicity manifested as pigmentary retinopathy, especially at higher doses, while the mechanism remains unelucidated.

The pharmacokinetics differences between the two enantiomers of CQ and HCQ have been documented both in animals and human.^42, 56, 58–59^ In a study of hydroxychloroquine distribution in rabbits, it was found that the *R*-enantiomer was stereoselectively accumulated in the ocular tissue with an *R/S*-HCQ ratio of 60/40 after the administration of the racemates. However, with an opposite ratio of 40:60 for the total plasma concentration, implying a possibility for a more significant contribution of the *R*-isomer for the ocular toxicity.^60^ The results were consistent with the data of clinical PK study in humans, where the whole blood concentration of the *R*-enantiomer was much higher than its antipode, whereas the ratio of R/S concentrations of the chiral metabolites was less than 1. Besides, the renal clearance for the *S*-enantiomer was twice greater than the *R*-enantiomer, and a shorter elimination half-life was observed with the *S*-enantiomer.^42, 56, 58–59^ *In vitro* assays using human plasma indicated that *R*-and *S*-HCQ enantiomers were 37% and 64% protein-bound, respectively, implying *R*-HCQ would have a higher volume of distribution than *S*-HCQ. These results indicated a faster renal excretion and/or more accentuated liver metabolism, and lower volume of distribution of the *S*-enantiomer, as compared to those of the *R*-enantiomer.^61^ Thus, a stereoselective elimination for the S-isomer and a stereoselective protein for the R-isomer were considered as attribution to the differences in disposition and distribution into tissues such as ocular compartment.

The pharmacological effects were also proven to be different between the two enantiomers of CQ and HCQ for various applications. The difference in their anti-malarial activities was reported in 1994.^44^ It was noted that by scientists at Sterling Winthrop Inc. that *S*-HCQ was approximately 70% more active than *R*-HCQ in the Rat Pleurisy Macrophage Model.^62^ Later on, *S*- and *R-*HCQ were found to suppress HIV-1 replication to a similar extent and dose proportionally, and were no more toxic than racemic HCQ.^63^ Since HCQ’s anti-malarial and antiviral effects had been purported to originate from its ability to increase endosomal pH, the authors demonstrated that both enantiomers of HCQ increased endosomal pH to similar levels in a dose proportionally manner.

Unlike the much-explored pharmacokinetics and pharmacological effects of the racemates, limited toxicity information has shown that *S*-chloroquine seems to be less toxic than the *R*-chloroquine.^43^ No further animal studies or clinical evidence was presented about the pharmacological activities of each enantiomer in the studies of malaria or autoimmune diseases.

The efficacy and safety of CQ and/or HCQ for treating COVD-19 need to be validated by larger and more definitive clinical trials. The complicated effects of ages, doses,^64^ disease states, prior treatments, underlying physiological conditions of enrolled patients should be taking into consideration, for all of which can cause bias in the trial outcomes. However, running trials with a racemic drug represents one of the most significant variances, i.e. with a drug substance of only 50% potency of the preferred isomer, and will unnecessarily complicate the delineation of clinical outcomes contributed by each enantiomer. Fortunately, racemic HCQ is readily available, with a worldwide production volume exceeding 300 metric tons and growing. Separation of a racemic mixture into pure enantiomer forms is industrially feasible, and asymmetric synthesis is also very promising as long-term solutions. There seems to be no excuse to continue running a trial using racemic CQ or HCQ other than ready availability.

### Different properties of CQ, HCQ, and their enantiomers against SARS-CoV-2, hERG, QT, and M^pro^

With this backdrop, we separated both racemic mixtures of CQ and HCQ by chiral HPLC, and tested the antiviral effects of *R*- and *S*-CQ and HCQ, as well as the racemic CQ, HCQ, and azithromycin in parallel. The result is illustrated in Figures 3 and 4. The most significant finding from this study is that both *S*-CQ and *S*-HCQ exhibited more pronounced activities than their respective *R*-enantiomer in the assay against SARS-CoV-2 *in vitro.* Specifically, the result with *S*-HCQ demonstrated significant enantioselectivity with 60 % more efficiency than *R*-HCQ. The IC_50_ of *S*-enantiomer of CQ and HCQ were 1.761 μM and 1.444 μM, respectively, compared with their *R*-enantiomers with a respective IC_50_ value of 1.975 μM and 2.445 μM. The IC 50 value of the racemic mixtures of CQ and HCQ was 1.801 μM and 1.752 μM, respectively, which fell between the two enantiomers, as expected. The IC_50_ of the racemic mixture of both CQ and HCQ indicated that, at the testing conditions, the *S*-isomers were the major contributors to the antiviral activities. It implies that the *R*-isomers were either coming for the ride, being minor contributors to the antiviral activities, providing hitherto unknown benefits such as cytokine storm management, or contributing to untoward side effects.

In addition to the antiviral efficiency of chiral CQ and HCQ against COVID-19, their cardiac toxicity as one major concern in clinical trials has been brought back to the spotlight after decades of using for malaria and autoimmune diseases. It has been documented that CQ or/and HCQ may induce QT prolongation and cardiomyopathy with a longer duration of treatment. However, it is not clear that these heart conduction disorders are associated with a specific enantiomer or simply from both forms, which would need a large prospective study for further assessment. We, for the first time, have demonstrated the prospect for superior safety profile concerning cardiovascular complications of *S-*HCQ and *S-*CQ over their respective *R-*isomers. Though the IC_50_ of racemic HCQ is over 10 μM that a concentration most drugs would accept for hERG activity in the early drug development stage, the extremely long half-life and the large volume of distribution of HCQ must be considered as well. A low/moderate hERG potency might still be a problem if the drug accumulated in the system for a long time, for cases as HCQ. *S-*HCQ showed an incredibly high IC_50_ (greater than 20 μM) at hERG inhibition, suggesting a much less chance of cardiac toxicity from the enantiomerically pure *S*-HCQ, as compared with racemic CQ and HCQ. Most importantly, *S*-HCQ alone did not induce QT prolongation *in vivo.* On the other hand, the combination of *S*-HCQ and azithromycin increased QT prolongation. Based on the analysis of the U.S. FDA’s Adverse Event Reporting System (FAERS) data by Amir Sarayani et al., ^65^ AZM treatment alone was significantly associated with TdP/QT prolongation. Our data suggest that the cardiotoxicity may be induced by the use of AZM at a certain level from the observation of significant prolongation of QT interval, but not by *S*-HCQ alone, which might be one explanation of the cardiac problems reported in the clinical trials applying HCQ and AZM therapy. Therefore, these results may lead to the speculation that *S*-HCQ alone has little effect on QT prolongation *in vivo*, while the combination with AZM may cause QT prolongation due to AZM related cardiotoxicity. Taking into account of aforementioned data on the more potent viral inhibition, it is expected that the therapeutic window could be effectively expanded by using pure *S*-HCQ substantially free of the *R*-isomer.

Though CQ and HCQ have been prescribed for decades, the mechanism of action still remains elusive. From the standpoint of the antiviral activity of CQ and HCQ against SARS-CoV-2, several mechanisms have been postulated. Due to their week base property, CQ and HCQ favorably accumulate in acidic compartments, such as endosomes and the Golgi apparatus; as a result, the intracellular pH increased. This effect would reduce the binding of ACE2 and the viral spike protein.^45^ In addition, as CQ and HCQ accumulate in other acidic organelles or inflamed tissue, it would lead to the interference of cytokine production and introduce immunomodulatory effects,^57^ which may be beneficial for the COVID-19 patients on the moderate or severe conditions, especially for the ones battling cytokine storm. Besides these modes of action proposed, many other proteins, such as RdRp, PLP, and M^pro^, also attract scientists and doctors’ attention as alternative drug targets. ^52, 66–69^ M^Pro^ was shown to be the putative viral protease responsible for unpacking the viral machiner for replications within host cells. As a preliminary attempt to understand the antiviral mechanisms of CQ and HCQ, which can be multiple, we subjected the enantiomers of CQ and HCQ to enzyme inhibition studies. Surprisingly, the *S-*enantiomers of both CQ and HCQ exhibited a much better inhibition towards M^pro^ than the *R-*enantiomers and their racemates. The IC_50_ values of HCQ series are nearly half of that in the CQ series, accordingly. Among all 6 tested compounds, *S-* HCQ showed the best potency with an IC_50_ value of 2.47 μM.

Since the mechanism of HCQ’s antiviral effect remains to be elucidated, we remain cautious at explaining the synergies between the antiviral and anti-inflammatory activities without enantioselectivity data for the latter.

## CONCLUSION

In summary, we have demonstrated the enantioselective antiviral activities of CQ and HCQ against SARS-CoV-2, hERG inhibition, and M^pro^ activity *in vitro*, as well as QT prolongation of *S*-HCQ *in vivo*. The *S*-enantiomers of CQ and HCQ proved more active than their corresponding *R*-enantiomers, suggesting the *in vitro* antiviral activities mainly derived from the *S*-isomers. We also showed an unequivocally pronounced difference in hERG activity of enantiomers of CQ and HCQ, indicating the best safety profile might be obtained from pure *S*-HCQ (hERG IC_50_ > 20 μM) and QT prolongation was not observed with *S*-HCQ alone. Therefore, it is reasonable to expect that the drug dose can be decreased to half of that of the racemates for a similar antiviral effect by removing the *R*-isomer. As a result, the untoward side effects attributed to the presence of the *R*-isomer, such as QT prolongation or cardiac toxicity as well as retinal damage, might be reduced to acceptable levels by applying only *S*-HCQ as an enantiomerically pure drug. Ambiguous or unfavorable outcome obtained with racemic HCQ from the clinical trials might be clarified with pure *S*-HCQ. In addition, we have provided evidence that M^pro^ could be a possible target for CQ and HCQ for treating COVID-19, and that M^pro^ exhibited a better binding affinity for the S- than the *R*-enantiomers. Racemate CQ or HCQ would be less preferred for COVID-19 applications unless shown to be so by further pharmacological testing. Taken these new data together with previous pharmacological and pharmacokinetic studies, we strongly recommend further human clinical studies be conducted with pure *S*-HCQ as the active pharmaceutical ingredient, constituting an expedient and highly probable way to materially increase the therapeutic index of HCQ for treating COVID-9.

## METHODS

### Material: Virus and cells

African green monkey kidney Vero E6 cell line was obtained from American Type Culture Collection (ATCC, cat no. 1586) and maintained in Dulbecco’s Modified Eagle’s medium (DMEM, Gibco) supplemented with 10% fetal bovine serum (FBS) at 37 °C incubator with 5% CO_2_. A clinical isolate SARS-CoV-2 virus (Genebank accession no. MT123290.1) was propagated in Vero E6 cells and viral titer was determined by plaque assay. All the infection experiments were performed in a Biosafety Level 3 (BLS 3) laboratory.

### Material: antivirals

The racemic chloroquine diphosphate (CAS NO. 50-63-5) and hydroxychloroquine sulfate (CAS NO. 747-36-4), purchased from Bidepharm, were converted into free racemic chloroquine and hydroxychloroquine, respectively, under basic conditions. The enantiomers of the racemic mixtures were separated by preparative chiral chromatography as free-bases with a Shimadzu LC-20AD HPLC equipment: chloroquine or hydroxychloroquine was dissolved in isocratic *n*-hexane/*iso*propanol/diethylamine with a ratio of 85:15:0.1 (v/v/v), and the loading needed for CQ and HCQ was 24.0 mg/mL and 23.5 mg/mL, respectively. The resulting solution was loaded onto a CHIRALPAK AY-H (AYH0CE-VC001) chiral column, which was then eluted with the same solvent system. The preparation conditions were as follows: the flow rate was 1.0 mL/min, the detection wavelength was UV 254 nm, and the temperature was 35 °C. The fractions of each enantiomer were collected and combined. The pure optical isomer was obtained by removing the solvent under reduced pressure with a rotary evaporator. The ee value of each enantiomer of both chloroquine and hydroxychloroquine was higher than 98%. The spectral data were identical to those reported in the literature. ^70–71^ These free-base enantiomers were converted to the enantiomerically pure diphosphate salt and sulfate salt, respectively, as described in the experimental section.

### SARS-CoV-2 antiviral assay and Immunofluorescence microscopy

Antiviral assays against SARS-CoV-2 *in vitro* were performed in a biosafety level-3 laboratory (BSL-3) at the State Key Laboratory of Respiratory Disease, Guangzhou Medical University, Guangzhou Customs District Technology Center.

Log phase Vero E6 cells were plated overnight in 96-well cell culture plate (Greiner, Cat no.655090) and then pretreated with the indicated concentration of compounds for 1 h (each concentration had three replicates). SARS-CoV-2, with a multiplicity of infection (MOI), of 0.05, was subsequently added. After 1 h, the medium was replaced with a fresh drug-containing one. 24 hours later, the cells were fixed with 4% paraformaldehyde and permeabilized with 0.2% Triton X 100. After blocking with 5% bovine serum albumin (BSA) at room temperature for 1 h, the rabbit polyclonal antibody against SARS Coronavirus Nucleoprotein (NP) (1:1000) as the primary antibody was used to incubate the cells, and then Alexa 488 labeled secondary antibody (Donkey Anti-Rabbit IgG;1:500; Jackson) was added. The nuclei were stained with DAPI (Sigma). The plate was scanned using Celigo Image Cytometer and fluorescence microscopy images were taken. The inhibition rate was calculated using the mean intensity of a green channel versus a DAPI channel. The IC_50_ value for each compound was calculated using GraphPad Prism software (version 7.0) using a non-linear regression equation.

### hERG inhibition assay

#### Cell culture and compound preparation

hERG-CHO cells were cultured in T175 flasks to a maximum 70-80% confluence at 37 °C and 5% CO_2_ incubator. Remove Culture Media (F-12 medium supplemented with 10% fetal bovine serum,100 g/mL G418 and 100 g/mL Hygromycin B) and wash with 7 mL PBS. Followed by dissociation with 3 mL Detachine reagent in 37 °C incubator for about 3 min, and replace with 7 mL Serum-Free Media to gently resuspend cells by using pipet up and down for several times. Finally, harvest cells by 800 rpm centrifuge and adjust to 2-5 million/mL cells for automated Qpatch 16X experiments. Before the assay, compounds 20 mM DMSO stock solutions were diluted with an external solution to the desired concentrations. The DMSO concentration is 0.2%, which has no effects on hERG currents.

### Automated QPatch recording

The hERG current recording in the whole-cell patch-clamp configuration on QPatch 16X applied single-hole QPlate. The cells were voltage-clamped at a holding potential of −80 mV. hERG current was activated by depolarizing at +20 mV for 5 second, after which the current was taken back to - 50 mV for 5 sec to remove the inactivation and observe the deactivating tail current. The peak size of tail current was used to quantify hERG current amplitude. Raw data have been included statistics as membrane resistance Rm>100 MΩ and tail current amplitude > 300 pA. External solution as followed (in mM, pH was adjusted to 7.4 by NaOH): 140 NaCl, 5 KCl, 1 CaCl_2_, 1.25 MgCl_2_, 10 Glucose, 10 HEPES. Internal solution as followed (in mM, pH was adjusted to 7.2 by NaOH): 140 KCl, 1 CaCl_2_, 1 MgCl_2_, 10 HEPES, 10 EGTA and 4Na_2_-ATP.

### QT prolongation assay in vivo

Specific pathogen-free (SPF) Hartley guinea pigs, weighing about 300 g were randomly divided into three groups with 8 in each group: control group (Con), *S*-hydroxychloroquine sulfate group (*S*-HCQ) and *S*-hydroxychloroquine sulfate + Azimycin group (*S*-HCQ + AZM). The Con group was received normal saline. The *S*-HCQ group was received 30.84 mg/kg *S*-HCQ on the first day, and 15.42 mg/kg *S*-HCQ on the following five days, once a day by gavage. The *S*-HCQ + AZM group was received *S*-HCQ (30.8 mg/kg on day1 followed by 15.4 mg/kg per day, the next five days), and 38.54 mg/kg AZM the first day and 19.27 mg/kg from day 2 through day 5. The dosage of guinea pig was adjusted and converted according to the clinical dosage of human (*45*). Electrocardiogram (ECG) of guinea pigs were measured before, on the 3rd and 6th day of administration.

The Guinea pigs were anesthetized with 2 % isoflurane and placed in their supine position on a heating pad. The surface lead II electrocardiogram (ECG) was obtained from the limb electrodes. ECG was monitored with a PowerLab 4/35 physiological monitor (AD Instruments Inc., United States), and analyzed with LabChart Pro Software (AD Instruments Inc., United States). The QT and RR intervals of ECG were measured, and the corrected QT interval (QTc) was calculated by Van de Water formula (*46*). All Guinea pigs were bred and maintained in a SPF, AAALAC-accredited facility of the Guangdong Laboratory Animals Monitoring Institute, Guangzhou, China. The experiments were approved by the Institutional Animal Care and Use Committee (No. IACUC2020125).

### M^pro^ evaluation

The pGEX4T1-Mpro plasmid was constructed (AtaGenix, Wuhan) and transfected into the E. coli strain BL21 (CodonPlus, Stratagene). A GST-tagged protein was purified by GST-glutathione affinity chromatography and cleaved with thrombin. The purity of the recombinant protein was greater than 95%, as assessed by SDS–PAGE. The activity of M^pro^ was measured by continuous kinetic assays, using an identical fluorogenic substrate MCA-AVLQSGFR-Lys(Dnp)-Lys-NH2 (Apetide Co., Ltd, Shanghai, China). The fluorescence intensity was monitored with a Multifunctional Enzyme Marker (SpectraMax^®^i3x, Molecular Divices, U.S.A.) using wavelengths of 320 and 405 nm for excitation and emission, respectively. The experiments were performed in a 100 μL reaction system with a buffer consisting of 50 mM Tris-HCl (pH 7.3), 1 mM EDTA. To determine the IC_50_ values, the test compounds were diluted in 100% DMSO to indicated concentration. The solution containing M^pro^ (final concentration of 500 nM) was dispensed into black 96-well plates with glass-bottom (Jing’an, Shanghai, China) and was incubated with 1 μL stock solution of test compounds at room temperature for 10 min. The reaction was initiated by adding substrate (final concentration of 20 μM). Fluorescence was monitored at 1 point per 45 s. Initial velocities were calculated by fitting the linear portion of the curves (the first 5 min of the progress curves) to a straight line using the program SoftMax Pro, and were converted to enzyme activity (substrate cleaved).

## EXPERIMENTAL

### Preparation of racemic chloroquine as a free-base (CAS NO. 54-05-7)

The purchased chloroquine diphosphate (13 g, 25.2 mmol) was dissolved in water (100 mL), followed by the addition of a 12% NaOH solution (50 mL) dropwise at 0 °C. After stirring for 0.5 h, EtOAc (25 mL) was added and the mixture kept stirring for another 0.5 h at rt. The reaction mixture was extracted with EtOAc (3 x 100 mL). The organic phase was combined and washed with brine, then dried with anhydrous Na_2_SO_4_. The organic solvent was removed by a rotary evaporator under reduced pressure to obtain a yellow residue as the free chloroquine (7.6 g, 23.8 mmol, 94.3 % yield). The spectral data were identical to the literature. ^72–73^

### Preparation of racemic hydroxychloroquine as a free-base (CAS NO. 118-42-3)

The purchased hydroxychloroquine sulfate (10.9 g, 27.4 mmol) was dissolved in water (75 mL), followed by the addition of a 12% NaOH solution (50 mL) dropwise at 0 °C. After stirring for 0.5 h, EtOAc (25 mL) was added and the mixture kept stirring for another 0.5 h at rt. The reaction mixture was extracted with EtOAc (3 × 100 mL). The organic phase was combined and washed with brine, then dried with anhydrous Na_2_SO_4_. The organic solvent was removed by a rotary evaporator under reduced pressure to obtain a yellow residue as the free hydroxychloroquine (7.7 g, 22.9 mmol, 83.7 % yield). The spectral data were identical to the literature.^74^

### Preparation of the enantiomers of chloroquine and hydroxychloroquine by chiral HPLC

The chiral high-pressure liquid chromatography (HPLC) method was used to separate the enantiomers of the racemic mixture as free-bases by with a Shimadzu LC-20AD HPLC equipment. Chloroquine or hydroxychloroquine was dissolved in isocratic *n*-hexane/*iso*propanol/diethylamine with a ratio of 85:15:0.1 (v/v/v), and the loading needed for CQ and HCQ was 24.0 mg/mL and 23.5 mg/mL, respectively. The resulting solution was loaded onto a CHIRALPAK AY-H (AYH0CE-VC001) chiral column, which was then eluted with the same solvent system. The preparation conditions were as follows: the flow rate was 1.0 mL/min, the detection wavelength was UV 254 nm, and the temperature was 35 °C. The fractions of each enantiomer were collected and combined. The pure optical isomer was obtained by removing the solvent under reduced pressure with a rotary evaporator. The ee of each enantiomer of both chloroquine and hydroxychloroquine was higher than 98 %. The spectral data were identical to the literature. ^70–71^

### General conversion from enantiomerically pure chloroquine as a free-base to its corresponding salt

*R*-chloroquine or *S*-chloroquine (640 mg, 2.0 mmol) was dissolved in EtOH (4 mL) and heated to reflux, followed by the addition of 85% phosphoric acid (0.25 mL) for 2 h. A large amount of white solid precipitated out. The reaction mixture was cooled to rt and filtered. The resulting solid was washed with EtOH (3 x 1 mL) to obtain the product as (*R*)-chloroquine diphosphate or (*S*)-chloroquine diphosphate.

### (4R)-4-N-(7-chloroquinolin-4-yl)-1-N,1-N-diethylpentane-1,4-diamine; phosphoric acid (R-chloroquine diphosphate salt)

The compound obtained was a white solid,868 mg, 1.68 mmol, 84.1 % yield, [α]_D_^26.8^ = −74.4 (*c* = 0.5, H_2_O). ^1^H NMR (400 MHz, D_2_O) δ 8.22 (d, *J* = 7.3 Hz, 1H), 8.06 (d, *J* = 9.1 Hz, 1H), 7.62 (d, *J* = 2.1 Hz, 1H), 7.46 (dd, *J* = 9.1, 2.1 Hz, 1H), 6.79 (d, *J* = 7.3 Hz, 1H), 4.09 (d, *J* = 5.9 Hz, 1H), 3.24 – 3.00 (m, 6H), 1.93 – 1.68 (m, 4H), 1.40 (d, *J* = 6.5 Hz, 3H), 1.22 (td, *J* = 7.3, 1.9 Hz, 6H); ^13^C NMR (101 MHz, D_2_O) δ 155.17, 142.10, 139.11, 137.88, 127.13, 123.92, 118.81, 114.95, 98.52, 51.13, 49.45, 47.22, 31.85, 20.21, 18.71, 8.09.

### (4S)-4-N-(7-chloroquinolin-4-yl)-1-N,1-N-diethylpentane-1,4-diamine; phosphoric acid (S-chloroquine diphosphate salt)

The compound obtained was a white solid, 887 mg, 1.72 mmol, 86.0 % yield, [α]_D_^27.8^ = 79.7 (*c* = 0.5, H_2_O). ^1^H NMR (400 MHz, D_2_O) δ 8.27 (d, *J* = 7.3 Hz, 1H), 8.16 (d, *J* = 9.1 Hz, 1H), 7.76 (d, *J* = 2.0 Hz, 1H), 7.57 (dd, *J* = 9.1, 2.1 Hz, 1H), 6.84 (d, *J* = 7.3 Hz, 1H), 4.23 – 3.97 (m, 1H), 3.29 – 3.10 (m, 6H), 1.90 – 1.73 (m, 4H), 1.41 (d, *J* = 6.5 Hz, 3H), 1.24 (td, *J* = 7.3, 1.9 Hz, 6H); ^13^C NMR (101 MHz, D_2_O) δ 155.39, 142.13, 139.23, 138.10, 127.23, 124.02, 119.00, 115.17, 98.48, 51.13, 49.43, 47.30, 31.88, 20.21, 18.73, 8.10.

### General conversion from enantiomerically pure hydroxychloroquine as a free-base to its corresponding salt

*R*-hydroxychloroquine or *S*-hydroxychloroquine, 700 mg, 2.1 mmol) was dissolved in EtOH (2 mL) and heated to 60 °C, followed by addition of 80% phosphoric acid (188 mg) for 1 h. The reaction mixture was cooled to −20 °C, a large amount of white solid precipitated out and the solid was filtered. The resulting solid was washed with cold EtOH (3 × 1 mL) to obtain the product as (*R*)-hydroxychloroquine sulfate or (*S*)-hydroxychloroquine sulfate.

### 2-[[(4R)-4-[(7-chloroquinolin-4-yl)amino]pentyl]-ethylamino]ethanol; sulfuric acid (R-hydroxychloroquine sulfate salt)

The compound obtained was a white solid, 745 mg, 1.71 mmol, 81.7 % yield, [α]_D_^26.1^ = −107.75 (*c* = 0.32, H_2_O). ^1^H NMR (400 MHz, D_2_O) δ 8.28 (d, *J* = 7.3 Hz, 1H), 8.16 (d, *J* = 9.1 Hz, 1H), 7.75 (d, *J* = 2.1 Hz, 1H), 7.56 (dd, *J* = 9.1, 2.1 Hz, 1H), 6.84 (d, *J* = 7.3 Hz, 1H), 4.14 (d, *J* = 6.5 Hz, 1H), 3.88 (t, *J* = 4.8 Hz, 2H), 3.34 – 3.16 (m, 6H), 1.96 – 1.71 (m, 4H), 1.42 (d, *J* = 6.5 Hz, 3H), 1.28 (td, *J* = 7.3, 1.8 Hz, 3H); ^13^C NMR (101 MHz, D_2_O) δ 155.34, 142.15, 139.17, 138.05, 127.20, 124.06, 118.98, 115.14, 98.53, 55.26, 53.77, 52.02, 49.46, 48.34, 31.86, 20.00, 18.74, 7.85.

### 2-[[(4S)-4-[(7-chloroquinolin-4-yl)amino]pentyl]-ethylamino]ethanol; sulfuric acid (S-hydroxychloroquine sulfate salt)

The compound obtained was a white solid (805 mg, 1.85 mmol, 88.3 % yield, [α]_D_^26.8^ = 95.6 (*c* = 0.32, H_2_O). ^1^H NMR (400 MHz, D_2_O) δ 8.26 (d, *J* = 7.3 Hz, 1H), 8.13 (d, *J* = 9.1 Hz, 1H), 7.69 (d, *J* = 2.1 Hz, 1H), 7.52 (dd, *J* = 9.1, 2.1 Hz, 1H), 6.83 (d, *J* = 7.3 Hz, 1H), 4.12 (dd, *J* = 12.6, 6.3 Hz, 1H), 3.88 (t, *J* = 5.0 Hz, 2H), 3.37 – 3.21 (m, 6H), 1.92 – 1.78 (m, 4H), 1.42 (d, *J* = 6.5 Hz, 3H), 1.28 (td, *J* = 7.3, 2.0 Hz, 3H); ^13^C NMR (101 MHz, D_2_O) δ 155.26, 142.15, 139.12, 137.96, 127.15, 124.03, 118.90, 115.05, 98.54, 55.27, 53.77, 52.03, 49.47, 48.26, 31.84, 20.01, 18.73, 7.85.

## ACKNOWLEDGMENT

We thank Shiyi Chen, Dongfeng Gu, Yu Zhang, Albert S. C. Chan and Erwei Song for their scientific discussion and helpful comments.

## Funding

We are grateful for financial support from Shenzhen Science and Technology Innovation Committee (ZDSYS20190902093215877) (X.Z.), Shenzhen Bay Laboratory (SZBL2019062801006) (X.Z.) and Guangdong Provincial Key Laboratory of Catalysis (No. 2020B121201002) (X.Z.) Southern University of Science and Technology. This work is also supported by the grants from The National Key Research and Development Program of China (2018YFC1200100) (J.Z.), National Science and Technology Major Project (2018ZX10301403) (J.Z.), the emergency grants for prevention and control of SARS-CoV-2 of Ministry of Science and Technology (2020YFC0841400) (J.Z.), Guangdong province (2020B111108001, 2018B020207013) (J.Z.), Science Foundation of Guangzhou City (201904020023) (H.-B.L.), Guangdong Provincial Key Laboratory of Construction Foundation (2017B030314030) (H.-B.L.), Local Innovative and Research Teams Project of Guangdong Pearl River Talents Program (2017BT01Y093) (H.-B.L.), Sun Yat-Sen University and Zhejiang University special scientific research fund for COVID-19 prevention and control (H.-B.L.) as well as Guangdong Province Key Laboratory of Laboratory Animals (2017B030314171) (F.Y.). All animal experiments were approved by the Institutional Animal Care and Use Committee of Guangdong Laboratory Animals Monitoring Institute (No. IACUC2020125).

## Author contributions

X.Z. and T.Z. initiated the project; X.Z., T.Z., J.Z. H.-B.L, and F.Y. designed the project. G.L. and T.Z. wrote the manuscript. G.L., Y.L., and Y.S. performed chiral HPLC separations the syntheses of tested compounds; J.S. performed the antiviral activity experiments; Y.H. carried out the M^pro^ activity experiments and hERG assay. X.L. performed the QT prolongation assay. G.L., J.S., Y.H., and Y.L. analyzed the data. X.Z., J.Z., H.-B.L., and F.Y. supervised and supported the project. All authors reviewed and approved the manuscript.

## Competing interests

All authors have no potential conflict of interest.

## Main contact

Tony Y. Zhang and Xumu Zhang

## Supplementary Materials

Figures S1-S14

## Notes

### Competing Interest Statement

The authors have declared no competing interest.

### Summary of Updates

The updated version included new data: hERG inhibition and Mpro activity in vitro of CQ, HCQ and their enantiomers, as well as QT prolongation evaluation in vivo of S-HCQ.

## REFERENCES

1. Huang, C.; Wang, Y.; Li, X.; Ren, L.; Zhao, J.; Hu, Y.; Zhang, L.; Fan, G.; Xu, J.; Gu, X.; Cheng, Z.; Yu, T.; Xia, J.; Wei, Y.; Wu, W.; Xie, X.; Yin, W.; Li, H.; Liu, M.; Xiao, Y.; Gao, H.; Guo, L.; Xie, J.; Wang, G.; Jiang, R.; Gao, Z.; Jin, Q.; Wang, J.; Cao, B., Clinical features of patients infected with 2019 novel coronavirus in Wuhan, China. Lancet 2020, 395 10223, 497–506.

2. Wu, F.; Zhao, S.; Yu, B.; Chen, Y.-M.; Wang, W.; Song, Z.-G.; Hu, Y.; Tao, Z.-W.; Tian, J.-H.; Pei, Y.-Y.; Yuan, M.-L.; Zhang, Y.-L.; Dai, F.-H.; Liu, Y.; Wang, Q.-M.; Zheng, J.-J.; Xu, L.; Holmes, E. C.; Zhang, Y.-Z., A new coronavirus associated with human respiratory disease in China. Nature 2020, 579 7798, 265–269.

3. Holshue, M. L.; DeBolt, C.; Lindquist, S.; Lofy, K. H.; Wiesman, J.; Bruce, H.; Spitters, C.; Ericson, K.; Wilkerson, S.; Tural, A.; Diaz, G.; Cohn, A.; Fox, L.; Patel, A.; Gerber, S. I.; Kim, L.; Tong, S.; Lu, X.; Lindstrom, S.; Pallansch, M. A.; Weldon, W. C.; Biggs, H. M.; Uyeki, T. M.; Pillai, S. K., First Case of 2019 Novel Coronavirus in the United States. N. Engl. J. Med. 2020, 382 10, 929–936.

4. Phan, L. T.; Nguyen, T. V.; Luong, Q. C.; Nguyen, T. V.; Nguyen, H. T.; Le, H. Q.; Nguyen, T. T.; Cao, T. M.; Pham, Q. D., Importation and Human-to-Human Transmission of a Novel Coronavirus in Vietnam. N. Engl. J. Med. 2020, 382 9, 872–874.

5. Rothe, C.; Schunk, M.; Sothmann, P.; Bretzel, G.; Froeschl, G.; Wallrauch, C.; Zimmer, T.; Thiel, V.; Janke, C.; Guggemos, W.; Seilmaier, M.; Drosten, C.; Vollmar, P.; Zwirglmaier, K.; Zange, S.; Wölfel, R.; Hoelscher, M., Transmission of 2019 - nCoV Infection from an Asymptomatic Contact in Germany. N. Engl. J. Med. 2020, 382 10, 970–971.

6. Zhu, N.; Zhang, D.; Wang, W.; Li, X.; Yang, B.; Song, J.; Zhao, X.; Huang, B.; Shi, W.; Lu, R.; Niu, P.; Zhan, F.; Ma, X.; Wang, D.; Xu, W.; Wu, G.; Gao, G. F.; Tan, W., A Novel Coronavirus from Patients with Pneumonia in China, 2019. N. Engl. J. Med. 2020, 382 8, 727–733.

7. https://gisanddata.maps.arcgis.com/apps/opsdashboard/index.html#/bda7594740fd40299423467b48e9ecf6.

8. Gao, Y.; Yan, L.; Huang, Y.; Liu, F.; Zhao, Y.; Cao, L.; Wang, T.; Sun, Q.; Ming, Z.; Zhang, L.; Ge, J.; Zheng, L.; Zhang, Y.; Wang, H.; Zhu, Y.; Zhu, C.; Hu, T.; Hua, T.; Zhang, B.; Yang, X.; Li, J.; Yang, H.; Liu, Z.; Xu, W.; Guddat, L. W.; Wang, Q.; Lou, Z.; Rao, Z., Structure of RNA-dependent RNA polymerase from 2019-nCoV, a major antiviral drug target. bioRxiv 2020, 2020.03.16.993386.

9. Gordon, D. E.; Jang, G. M.; Bouhaddou, M.; Xu, J.; Obernier, K.; O’Meara, M. J.; Guo, J. Z.; Swaney, D. L.; Tummino, T. A.; Huettenhain, R.; Kaake, R. M.; Richards, A. L.; Tutuncuoglu, B.; Foussard, H.; Batra, J.; Haas, K.; Modak, M.; Kim, M.; Haas, P.; Polacco, B. J.; Braberg, H.; Fabius, J. M.; Eckhardt, M.; Soucheray, M.; Bennett, M. J.; Cakir, M.; McGregor, M. J.; Li, Q.; Naing, Z. Z. C.; Zhou, Y.; Peng, S.; Kirby, I. T.; Melnyk, J. E.; Chorba, J. S.; Lou, K.; Dai, S. A.; Shen, W.; Shi, Y.; Zhang, Z.; Barrio-Hernandez, I.; Memon, D.; Hernandez-Armenta, C.; Mathy, C. J. P.; Perica, T.; Pilla, K. B.; Ganesan, S. J.; Saltzberg, D. J.; Ramachandran, R.; Liu, X.; Rosenthal, S. B.; Calviello, L.; Venkataramanan, S.; Liboy-Lugo, J.; Lin, Y.; Wankowicz, S. A.; Bohn, M.; Sharp, P. P.; Trenker, R.; Young, J. M.; Cavero, D. A.; Hiatt, J.; Roth, T. L.; Rathore, U.; Subramanian, A.; Noack, J.; Hubert, M.; Roesch, F.; Vallet, T.; Meyer, B.; White, K. M.; Miorin, L.; Rosenberg, O. S.; Verba, K. A.; Agard, D.; Ott, M.; Emerman, M.; Ruggero, D.; García -Sastre, A.; Jura, N.; von Zastrow, M.; Taunton, J.; Ashworth, A.; Schwartz, O.; Vignuzzi, M.; d’Enfert, C.; Mukherjee, S.; Jacobson, M.; Malik, H. S.; Fujimori, D. G.; Ideker, T.; Craik, C. S.; Floor, S.; Fraser, J. S.; Gross, J.; Sali, A.; Kortemme, T.; Beltrao, P.; Shokat, K.; Shoichet, B. K.; Krogan, N. J., A SARS-CoV-2-Human Protein-Protein Interaction Map Reveals Drug Targets and Potential Drug-Repurposing. bioRxiv 2020, 2020.03.22.002386.

10. Zhou, Y.; Hou, Y.; Shen, J.; Huang, Y.; Martin, W.; Cheng, F., Network-based drug repurposing for novel coronavirus 2019-nCoV/SARS-CoV-2. Cell Discovery 2020, 6 1, 14.

11. Wang, M.; Cao, R.; Zhang, L.; Yang, X.; Liu, J.; Xu, M.; Shi, Z.; Hu, Z.; Zhong, W.; Xiao, G., Remdesivir and chloroquine effectively inhibit the recently emerged novel coronavirus (2019-nCoV) in vitro. Cell Res. 2020, 30 3, 269–271.

12. Huang, M.; Tang, T.; Pang, P.; Li, M.; Ma, R.; Lu, J.; Shu, J.; You, Y.; Chen, B.; Liang, J.; Hong, Z.; Chen, H.; Kong, L.; Qin, D.; Pei, D.; Xia, J.; Jiang, S.; Shan, H., Treating COVID-19 with Chloroquine. J. Mol. Cell. Biol. 2020.

13. Yao, X.; Ye, F.; Zhang, M.; Cui, C.; Huang, B.; Niu, P.; Liu, X.; Zhao, L.; Dong, E.; Song, C.; Zhan, S.; Lu, R.; Li, H.; Tan, W.; Liu, D., In Vitro Antiviral Activity and Projection of Optimized Dosing Design of Hydroxychloroquine for the Treatment of Severe Acute Respiratory Syndrome Coronavirus 2 (SARS-CoV-2). Clin. Infect. Dis. 2020.

14. Deng, L.; Li, C.; Zeng, Q.; Liu, X.; Li, X.; Zhang, H.; Hong, Z.; Xia, J., Arbidol combined with LPV/r versus LPV/r alone against Corona Virus Disease 2019: A retrospective cohort study. J. Infect. 2020.

15. Khalili, J. S.; Zhu, H.; Mak, A.; Yan, Y.; Zhu, Y., Novel coronavirus treatment with ribavirin: Groundwork for evaluation concerning COVID-19. J. Med. Virol. 2020.

16. Elfiky, A. A., Anti-HCV, nucleotide inhibitors, repurposing against COVID-19. Life Sci. 2020, 248, 117477.

17. Elfiky, A. A., Ribavirin, Remdesivir, Sofosbuvir, Galidesivir, and Tenofovir against SARS-CoV-2 RNA dependent RNA polymerase (RdRp): A molecular docking study. Life Sci. 2020, 117592.

18. Wang, Z.; Chen, X.; Lu, Y.; Chen, F.; Zhang, W., Clinical characteristics and therapeutic procedure for four cases with 2019 novel coronavirus pneumonia receiving combined Chinese and Western medicine treatment. Biosci. Trends 2020, 14 1, 64–68.

19. Liu, F.; Xu, A.; Zhang, Y.; Xuan, W.; Yan, T.; Pan, K.; Yu, W.; Zhang, J., Patients of COVID-19 may benefit from sustained lopinavir-combined regimen and the increase of eosinophil may predict the outcome of COVID-19 progression. Int. J. Infect. Dis. 2020.

20. Cao, B.; Wang, Y.; Wen, D.; Liu, W.; Wang, J.; Fan, G.; Ruan, L.; Song, B.; Cai, Y.; Wei, M.; Li, X.; Xia, J.; Chen, N.; Xiang, J.; Yu, T.; Bai, T.; Xie, X.; Zhang, L.; Li, C.; Yuan, Y.; Chen, H.; Li, H.; Huang, H.; Tu, S.; Gong, F.; Liu, Y.; Wei, Y.; Dong, C.; Zhou, F.; Gu, X.; Xu, J.; Liu, Z.; Zhang, Y.; Li, H.; Shang, L.; Wang, K.; Li, K.; Zhou, X.; Dong, X.; Qu, Z.; Lu, S.; Hu, X.; Ruan, S.; Luo, S.; Wu, J.; Peng, L.; Cheng, F.; Pan, L.; Zou, J.; Jia, C.; Wang, J.; Liu, X.; Wang, S.; Wu, X.; Ge, Q.; He, J.; Zhan, H.; Qiu, F.; Guo, L.; Huang, C.; Jaki, T.; Hayden, F. G.; Horby, P. W.; Zhang, D.; Wang, C., A Trial of Lopinavir-Ritonavir in Adults Hospitalized with Severe Covid-19. N. Engl. J. Med. 2020.

21. Siegel, D.; Hui, H. C.; Doerffler, E.; Clarke, M. O.; Chun, K.; Zhang, L.; Neville, S.; Carra, E.; Lew, W.; Ross, B.; Wang, Q.; Wolfe, L.; Jordan, R.; Soloveva, V.; Knox, J.; Perry, J.; Perron, M.; Stray, K. M.; Barauskas, O.; Feng, J. Y.; Xu, Y.; Lee, G.; Rheingold, A. L.; Ray, A. S.; Bannister, R.; Strickley, R.; Swaminathan, S.; Lee, W. A.; Bavari, S.; Cihlar, T.; Lo, M. K.; Warren, T. K.; Mackman, R. L., Discovery and Synthesis of a Phosphoramidate Prodrug of a Pyrrolo[2,1-f][triazin-4-amino] Adenine C-Nucleoside (GS-5734) for the Treatment of Ebola and Emerging Viruses. J. Med. Chem. 2017, 60 5, 1648–1661.

22. Warren, T. K.; Jordan, R.; Lo, M. K.; Ray, A. S.; Mackman, R. L.; Soloveva, V.; Siegel, D.; Perron, M.; Bannister, R.; Hui, H. C.; Larson, N.; Strickley, R.; Wells, J.; Stuthman, K. S.; Van Tongeren, S. A.; Garza, N. L.; Donnelly, G.; Shurtleff, A. C.; Retterer, C. J.; Gharaibeh, D.; Zamani, R.; Kenny, T.; Eaton, B. P.; Grimes, E.; Welch, L. S.; Gomba, L.; Wilhelmsen, C. L.; Nichols, D. K.; Nuss, J. E.; Nagle, E. R.; Kugelman, J. R.; Palacios, G.; Doerffler, E.; Neville, S.; Carra, E.; Clarke, M. O.; Zhang, L.; Lew, W.; Ross, B.; Wang, Q.; Chun, K.; Wolfe, L.; Babusis, D.; Park, Y.; Stray, K. M.; Trancheva, I.; Feng, J. Y.; Barauskas, O.; Xu, Y.; Wong, P.; Braun, M. R.; Flint, M.; McMullan, L. K.; Chen, S. S.; Fearns, R.; Swaminathan, S.; Mayers, D. L.; Spiropoulou, C. F.; Lee, W. A.; Nichol, S. T.; Cihlar, T.; Bavari, S., Therapeutic efficacy of the small molecule GS-5734 against Ebola virus in rhesus monkeys. Nature 2016, 531 7594, 381–5.

23. Million, M.; Lagier, J. C.; Gautret, P.; Colson, P.; Fournier, P. E.; Amrane, S.; Hocquart, M.; Mailhe, M.; Esteves-Vieira, V.; Doudier, B.; Aubry, C.; Correard, F.; Giraud-Gatineau, A.; Roussel, Y.; Berenger, C.; Cassir, N.; Seng, P.; Zandotti, C.; Dhiver, C.; Ravaux, I.; Tomei, C.; Eldin, C.; Tissot-Dupont, H.; Honoré, S.; Stein, A.; Jacquier, A.; Deharo, J. C.; Chabrière, E.; Levasseur, A.; Fenollar, F.; Rolain, J. M.; Obadia, Y.; Brouqui, P.; Drancourt, M.; La Scola, B.; Parola, P.; Raoult, D., Early treatment of COVID-19 patients with hydroxychloroquine and azithromycin: A retrospective analysis of 1061 cases in Marseille, France. Travel Med. Infect. Dis. 2020, 101738.

24. Huang, M.; Li, M.; Xiao, F.; Pang, P.; Liang, J.; Tang, T.; Liu, S.; Chen, B.; Shu, J.; You, Y.; Li, Y.; Tang, M.; Zhou, J.; Jiang, G.; Xiang, J.; Hong, W.; He, S.; Wang, Z.; Feng, J.; Lin, C.; Ye, Y.; Wu, Z.; Li, Y.; Zhong, B.; Sun, R.; Hong, Z.; Liu, J.; Chen, H.; Wang, X.; Li, Z.; Pei, D.; Tian, L.; Xia, J.; Jiang, S.; Zhong, N.; Shan, H., Preliminary evidence from a multicenter prospective observational study of the safety and efficacy of chloroquine for the treatment of COVID-19. National Science Review 2020.

25. Gao, J.; Tian, Z.; Yang, X., Breakthrough: Chloroquine phosphate has shown apparent efficacy in treatment of COVID-19 associated pneumonia in clinical studies. Biosci. Trends 2020, 14 1, 72–73.

26. Colson, P.; Rolain, J.-M.; Lagier, J.-C.; Brouqui, P.; Raoult, D., Chloroquine and hydroxychloroquine as available weapons to fight COVID-19. Int. J. Antimicrob. Agents 2020, 55 4, 105932–105932.

27. Gautret, P.; Lagier, J. C.; Parola, P.; Hoang, V. T.; Meddeb, L.; Mailhe, M.; Doudier, B.; Courjon, J.; Giordanengo, V.; Vieira, V. E.; Tissot Dupont, H.; Honoré, S.; Colson, P.; Chabrière, E.; La Scola, B.; Rolain, J. M.; Brouqui, P.; Raoult, D., Hydroxychloroquine and azithromycin as a treatment of COVID-19: results of an open-label non-randomized clinical trial. Int. J. Antimicrob. Agents 2020, 56 (1), 105949.

28. Gautret, P.; Lagier, J. C.; Parola, P.; Hoang, V. T.; Meddeb, L.; Sevestre, J.; Mailhe, M.; Doudier, B.; Aubry, C.; Amrane, S.; Seng, P.; Hocquart, M.; Eldin, C.; Finance, J.; Vieira, V. E.; Tissot-Dupont, H. T.; Honoré, S.; Stein, A.; Million, M.; Colson, P.; La Scola, B.; Veit, V.; Jacquier, A.; Deharo, J. C.; Drancourt, M.; Fournier, P. E.; Rolain, J. M.; Brouqui, P.; Raoult, D., Clinical and microbiological effect of a combination of hydroxychloroquine and azithromycin in 80 COVID-19 patients with at least a six-day follow up: A pilot observational study. Travel Med. Infect. Dis. 2020, 34, 101663.

29. Million, M.; Lagier, J.-C.; Gautret, P.; Colson, P.; Fournier, P.-E.; Amrane, S.; Hocquart, M.; Mailhe, M.; Esteves-Vieira, V.; Doudier, B.; Aubry, C.; Correard, F.; Giraud-Gatineau, A.; Roussel, Y.; Berenger, C.; Cassir, N.; Seng, P.; Zandotti, C.; Dhiver, C.; Ravaux, I.; Tomei, C.; Eldin, C.; Tissot-Dupont, H.; Honoré, S.; Stein, A.; Jacquier, A.; Deharo, J.-C.; Chabrière, E.; Levasseur, A.; Fenollar, F.; Rolain, J.-M.; Obadia, Y.; Brouqui, P.; Drancourt, M.; La Scola, B.; Parola, P.; Raoult, D., Early treatment of COVID-19 patients with hydroxychloroquine and azithromycin: A retrospective analysis of 1061 cases in Marseille, France. Travel Med. Infect. Dis. 2020, 35, 101738–101738.

30. Lagier, J. C.; Million, M.; Gautret, P.; Colson, P.; Cortaredona, S.; Giraud-Gatineau, A.; Honoré, S.; Gaubert, J. Y.; Fournier, P. E.; Tissot - Dupont, H.; Chabrière, E.; Stein, A.; Deharo, J. C.; Fenollar, F.; Rolain, J. M.; Obadia, Y.; Jacquier, A.; La Scola, B.; Brouqui, P.; Drancourt, M.; Parola, P.; Raoult, D., Outcomes of 3,737 COVID-19 patients treated with hydroxychloroquine/azithromycin and other regimens in Marseille, France: A retrospective analysis. Travel Med. Infect. Dis. 2020, 101791.

31. d’Arminio Monforte, A.; Tavelli, A.; Bai, F.; Marchetti, G.; Cozzi-Lepri, A., Effectiveness of Hydroxychloroquine in COVID-19 disease: A done and dusted situation? International Journal of Infectious Diseases.

32. Magagnoli, J.; Narendran, S.; Pereira, F.; Cummings, T. H.; Hardin, J. W.; Sutton, S. S.; Ambati, J., Outcomes of Hydroxychloroquine Usage in United States Veterans Hospitalized with COVID-19. Med 2020.

33. Chorin, E.; Dai, M.; Shulman, E.; Wadhwani, L.; Bar-Cohen, R.; Barbhaiya, C.; Aizer, A.; Holmes, D.; Bernstein, S.; Spinelli, M.; Park, D. S.; Chinitz, L. A.; Jankelson, L., The QT interval in patients with COVID-19 treated with hydroxychloroquine and azithromycin. Nat. Med. 2020, 26 6, 808–809.

34. Rosenberg, E. S.; Dufort, E. M.; Udo, T.; Wilberschied, L. A.; Kumar, J.; Tesoriero, J.; Weinberg, P.; Kirkwood, J.; Muse, A.; DeHovitz, J.; Blog, D. S.; Hutton, B.; Holtgrave, D. R.; Zucker, H. A., Association of Treatment With Hydroxychloroquine or Azithromycin With In-Hospital Mortality in Patients With COVID-19 in New York State. JAMA 2020, 323 24, 2493–502.

35. Mehra, M. R.; Desai, S. S.; Ruschitzka, F.; Patel, A. N., RETRACTED: Hydroxychloroquine or chloroquine with or without a macrolide for treatment of COVID-19: a multinational registry analysis. Lancet (London, England) 2020, S0140-6736(20)31180-6.

36. Horby, P.; Mafham, M.; Linsell, L.; Bell, J. L.; Staplin, N.; Emberson, J. R.; Wiselka, M.; Ustianowski, A.; Elmahi, E.; Prudon, B.; Whitehouse, A.; Felton, T.; Williams, J.; Faccenda, J.; Underwood, J.; Baillie, J. K.; Chappell, L.; Faust, S. N.; Jaki, T.; Jeffery, K.; Lim, W. S.; Montgomery, A.; Rowan, K.; Tarning, J.; Watson, J. A.; White, N. J.; Juszczak, E.; Haynes, R.; Landray, M. J., Effect of Hydroxychloroquine in Hospitalized Patients with COVID-19: Preliminary results from a multi-centre, randomized, controlled trial. medRxiv 2020, 2020.07.15.20151852.

37. Hoffmann, M.; Mösbauer, K.; Hofmann-Winkler, H.; Kaul, A.; Kleine-Weber, H.; Krüger, N.; Gassen, N. C.; Müller, M. A.; Drosten, C.; Pöhlmann, S., Chloroquine does not inhibit infection of human lung cells with SARS-CoV-2. Nature 2020.

38. Plantone, D.; Koudriavtseva, T., Current and Future Use of Chloroquine and Hydroxychloroquine in Infectious, Immune, Neoplastic, and Neurological Diseases: A Mini-Review. Clin. Drug Investig. 2018, 38 (8), 653–671.

39. Surrey, A. R., 7-chloro-4-[5-(n-ethyl-n-2-hydroxyethylamino)-2-pentyl] aminoquinoline, its acid addition salts, and method of preparation. US Patent 2546658A 1951.

40. Chew, C. Y.; Mar, A.; Nikpour, M.; Saracino, A. M., Hydroxychloroquine in dermatology: New perspectives on an old drug. Australas. J. Dermatol. 2019.

41. Shipman, W. D.; Vernice, N. A.; Demetres, M.; Jorizzo, J. L., An update on the use of hydroxychloroquine in cutaneous lupus erythematosus: A systematic review. J. Am. Acad. Dermatol. 2020, 82 (3), 709–722.

42. Tett, S. E.; McLachlan, A. J.; Cutler, D. J.; Day, R. O., Pharmacokinetics and pharmacodynamics of hydroxychloroquine enantiomers in patients with rheumatoid arthritis receiving multiple doses of racemate. Chirality 1994, 6 (4), 355–9.

43. Tagoe, C. N.; Ofori-Adjei, D., Effects of chloroquine and its enantiomers on the development of rat embryos in vitro. Teratology 1995, 52 (3), 137–42.

44. Vera J. Stecher, W. F. M., (S)-(+)-hydroxychloroquine. US Patent 5314894A 1994.

45. Liu, J.; Cao, R.; Xu, M.; Wang, X.; Zhang, H.; Hu, H.; Li, Y.; Hu, Z.; Zhong, W.; Wang, M., Hydroxychloroquine, a less toxic derivative of chloroquine, is effective in inhibiting SARS-CoV-2 infection in vitro. Cell Discovery 2020, 6 (1), 16.

46. Arshad, S.; Kilgore, P.; Chaudhry, Z. S.; Jacobsen, G.; Wang, D. D.; Huitsing, K.; Brar, I.; Alangaden, G. J.; Ramesh, M. S.; McKinnon, J. E.; O’Neill, W.; Zervos, M., Treatment with Hydroxychloroquine, Azithromycin, and Combination in Patients Hospitalized with COVID-19. Int. J. Infect. Dis. 2020, 97, 396–403.

47. Giudicessi, J. R.; Noseworthy, P. A.; Friedman, P. A.; Ackerman, M. J., Urgent Guidance for Navigating and Circumventing the QTc-Prolonging and Torsadogenic Potential of Possible Pharmacotherapies for Coronavirus Disease 19 (COVID-19). Mayo Clin. Proc. 2020, 95 (6), 1213–1221.

48. Sanguinetti, M. C.; Tristani-Firouzi, M., hERG potassium channels and cardiac arrhythmia. Nature 2006, 440 (7083), 463–469.

49. Yao, X.; Anderson, D. L.; Ross, S. A.; Lang, D. G.; Desai, B. Z.; Cooper, D. C.; Wheelan, P.; McIntyre, M. S.; Bergquist, M. L.; MacKenzie, K. I.; Becherer, J. D.; Hashim, M. A., Predicting QT prolongation in humans during early drug development using hERG inhibition and an anaesthetized guinea-pig model. Br. J. Pharmacol. 2008, 154 (7), 1446–1456.

50. Cortegiani, A.; Ingoglia, G.; Ippolito, M.; Giarratano, A.; Einav, S., A systematic review on the efficacy and safety of chloroquine for the treatment of COVID-19. J. Crit. Care 2020, 57, 279–283.

51. Van de Water, A.; Verheyen, J.; Xhonneux, R.; Reneman, R. S., An improved method to correct the QT interval of the electrocardiogram for changes in heart rate. J. Pharmacol. Methods 1989, 22 (3), 207–217.

52. Li, Z.; Li, X.; Huang, Y.-Y.; Wu, Y.; Liu, R.; Zhou, L.; Lin, Y.; Wu, D.; Zhang, L.; Liu, H.; Xu, X.; Yu, K.; Zhang, Y.; Cui, J.; Zhan, C.-G.; Wang, X.; Luo, H.-B., Identify potent SARS-CoV-2 main protease inhibitors via accelerated free energy perturbation-based virtual screening of existing drugs. bioRxiv 2020, 2020.03.23.004580.

53. Tokunaga, E.; Yamamoto, T.; Ito, E.; Shibata, N., Understanding the Thalidomide Chirality in Biological Processes by the Self-disproportionation of Enantiomers. Sci. Rep. 2018, 8 (1), 17131.

54. McLachlan, A. J.; Tett, S. E.; Cutler, D. J., High-performance liquid chromatographic separation of the enantiomers of hydroxychloroquine and its major metabolites in biological fluids using an alpha 1-acid glycoprotein stationary phase. J. Chromatogr. 1991, 570 (1), 119–27.

55. Ibrahim, K. E.; Fell, A. F., Separation of chloroquine enantiomers by high-performance liquid chromatography. J. Pharm. Biomed. Anal. 1990, 8 (5), 449–452.

56. McLachlan, A. J.; Tett, S. E.; Cutler, D. J.; Day, R. O., Disposition of the enantiomers of hydroxychloroquine in patients with rheumatoid arthritis following multiple doses of the racemate. Br. J. Clin. Pharmacol. 1993, 36 (1), 78–81.

57. Schrezenmeier, E.; Dörner, T., Mechanisms of action of hydroxychloroquine and chloroquine: implications for rheumatology. Nat. Rev. Rheumatol. 2020, 16 (3), 155–166.

58. Ducharme, J.; Fieger, H.; Ducharme, M. P.; Khalil, S. K.; Wainer, I. W., Enantioselective disposition of hydroxychloroquine after a single oral dose of the racemate to healthy subjects. Br. J. Clin. Pharmacol. 1995, 40 (2), 127–133.

59. Furst, D. E., Pharmacokinetics of hydroxychloroquine and chloroquine during treatment of rheumatic diseases. Lupus 1996, 5 ((1_suppl)), 11–15.

60. Wainer, I. W.; Chen, J. C.; Parenteau, H.; Abdullah, S.; Ducharme, J.; Fieger, H.; Iredale, J., Distribution of the enantiomers of hydroxychloroquine and its metabolites in ocular tissues of the rabbit after oral administration of racemic-hydroxychloroquine. Chirality 1994, 6 (4), 347–54.

61. Midha, K. K.; Hubbard, J. W.; Rawson, M. J.; McKay, G.; Schwede, R., The roles of stereochemistry and partial areas in a parallel design study to assess the bioequivalence of two formulations of hydroxychloroquine: A drug with a very long half life. Eur. J. Pharm. Sci. 1996, 4 (5), 283–292.

62. Wei, Y.; Nygard, G. A.; Ellertson, S. L.; Khalil, S. K. W., Stereoselective disposition of hydroxychloroquine and its metabolites in rats. Chirality 1995, 7 (8), 598–604.

63. Chiang, G.; Sassaroli, M.; Louie, M.; Chen, H.; Stecher, V. J.; Sperber, K., Inhibition of HIV-1 replication by hydroxychloroquine: mechanism of action and comparison with zidovudine. Clin. Ther. 1996, 18 (6), 1080–1092.

64. Yao, X.; Ye, F.; Zhang, M.; Cui, C.; Huang, B.; Niu, P.; Liu, X.; Zhao, L.; Dong, E.; Song, C.; Zhan, S.; Lu, R.; Li, H.; Tan, W.; Liu, D., In Vitro Antiviral Activity and Projection of Optimized Dosing Design of Hydroxychloroquine for the Treatment of Severe Acute Respiratory Syndrome Coronavirus 2 (SARS-CoV-2). Clin. Infect. Dis. 2020.

65. Sarayani, A.; Cicali, B.; Henriksen, C. H.; Brown, J. D., Safety signals for QT prolongation or Torsades de Pointes associated with azithromycin with or without chloroquine or hydroxychloroquine. Research in Social and Administrative Pharmacy 2020.

66. Adedeji, A. O.; Sarafianos, S. G., Antiviral drugs specific for coronaviruses in preclinical development. Curr. Opin. Virol. 2014, 8, 45–53.

67. Liu, X.; Li, Z.; Liu, S.; Sun, J.; Chen, Z.; Jiang, M.; Zhang, Q.; Wei, Y.; Wang, X.; Huang, Y.-Y.; Shi, Y.; Xu, Y.; Xian, H.; Bai, F.; Ou, C.; Xiong, B.; Lew, A. M.; Cui, J.; Fang, R.; Huang, H.; Zhao, J.; Hong, X.; Zhang, Y.; Zhou, F.; Luo, H.-B., Potential therapeutic effects of dipyridamole in the severely ill patients with COVID-19. Acta Pharmaceutica Sinica B 2020.

68. Dai, W.; Zhang, B.; Su, H.; Li, J.; Zhao, Y.; Xie, X.; Jin, Z.; Liu, F.; Li, C.; Li, Y.; Bai, F.; Wang, H.; Cheng, X.; Cen, X.; Hu, S.; Yang, X.; Wang, J.; Liu, X.; Xiao, G.; Jiang, H.; Rao, Z.; Zhang, L.-K.; Xu, Y.; Yang, H.; Liu, H., Structure-based design of antiviral drug candidates targeting the SARS-CoV-2 main protease. Science 2020, eabb4489.

69. Jin, Z.; Du, X.; Xu, Y.; Deng, Y.; Liu, M.; Zhao, Y.; Zhang, B.; Li, X.; Zhang, L.; Peng, C.; Duan, Y.; Yu, J.; Wang, L.; Yang, K.; Liu, F.; Jiang, R.; Yang, X.; You, T.; Liu, X.; Yang, X.; Bai, F.; Liu, H.; Liu, X.; Guddat, L. W.; Xu, W.; Xiao, G.; Qin, C.; Shi, Z.; Jiang, H.; Rao, Z.; Yang, H., Structure of Mpro from SARS-CoV-2 and discovery of its inhibitors. Nature 2020.

70. Blaney, P. M.; Byard, S. J.; Carr, G.; Ellames, G. J.; Herbert, J. M.; Peace, J. E.; Smith, D. I.; Michne, W. F.; Sanner, M. S., A practical synthesis of the enantiomers of hydroxychloroquine. Tetrahedron: Asymmetry 1994, 5 (9), 1815–1822.

71. Sinha, M.; Dola, V.; Soni, A.; Agarwal, P.; Srivastava, K.; Haq, W.; Puri, S. K.; Katti, S., Synthesis of chiral chloroquine and its analogues as antimalarial agents. Bioorg. Med. Chem. 2014, 22.

72. Staderini, M.; Bolognesi, M. L.; Menéndez, J. C., Lewis Acid - Catalyzed Generation of C C and C N Bonds on π-Deficient Heterocyclic Substrates. Advanced Synthesis & Catalysis 2015, 357 (1), 185–195.

73. De, D.; Byers, L. D.; Krogstad, D. J., Antimalarials: Synthesis of 4-aminoquinolines that circumvent drug resistance in malaria parasites. J. Heterocycl. Chem. 1997, 34 (1), 315–320.

74. Yu, E.; Mangunuru, H. P. R.; Telang, N. S.; Kong, C. J.; Verghese, J.; Gilliland Iii, S. E.; Ahmad, S.; Dominey, R. N.; Gupton, B. F., High-yielding continuous-flow synthesis of antimalarial drug hydroxychloroquine. Beilstein J. Org. Chem. 2018, 14, 583–592.

