## Supporting information for "Enantiomers of Chloroquine and Hydroxychloroquine Exhibit Different Activities Against SARS-CoV-2 *in vitro*, Evidencing *S*-Hydroxychloroquine as a Potentially Superior Drug for COVID-19"

DO NOT COPY

### Table of Contents

Figure S1. [1]H NMR (400 MHz, $D_2O$) of Rac-CQ diphosphate. 3
Figure S2. [13]C NMR (101 MHz, $D_2O$) of Rac-CQ diphosphate. 3
Figure S3. [1]H NMR (400 MHz, $D_2O$) of Rac-HCQ sulfate. 4
Figure S4. [13]C NMR (101 MHz, $D_2O$) of Rac-HCQ sulfate. 4
Figure S5. [1]H NMR (400 MHz, $D_2O$) of R-CQ diphosphate. 5
Figure S7. [1]H NMR (400 MHz, $D_2O$) of S-CQ diphosphate. 6
Figure S8. [13]C NMR (101 MHz, $D_2O$) of S-CQ diphosphate. 6
Figure S9. [1]H NMR (400 MHz, $D_2O$) of R-HCQ sulfate. 7
Figure S10. [13]C NMR (101 MHz, $D_2O$) of R-HCQ sulfate. 7
Figure S11. [1]H NMR (400 MHz, $D_2O$) of S-HCQ sulfate. 8
Figure S12. [13]C NMR (101 MHz, $D_2O$) of S-HCQ sulfate. 8
Figure S13. Antiviral activities (second batch) 9
Figure S14. Antiviral activities (statistics) 10

DO NOT COPY

Figure S1. $^{1}H$ NMR (400 MHz, $D_2O$) of Rac-CQ diphosphate.

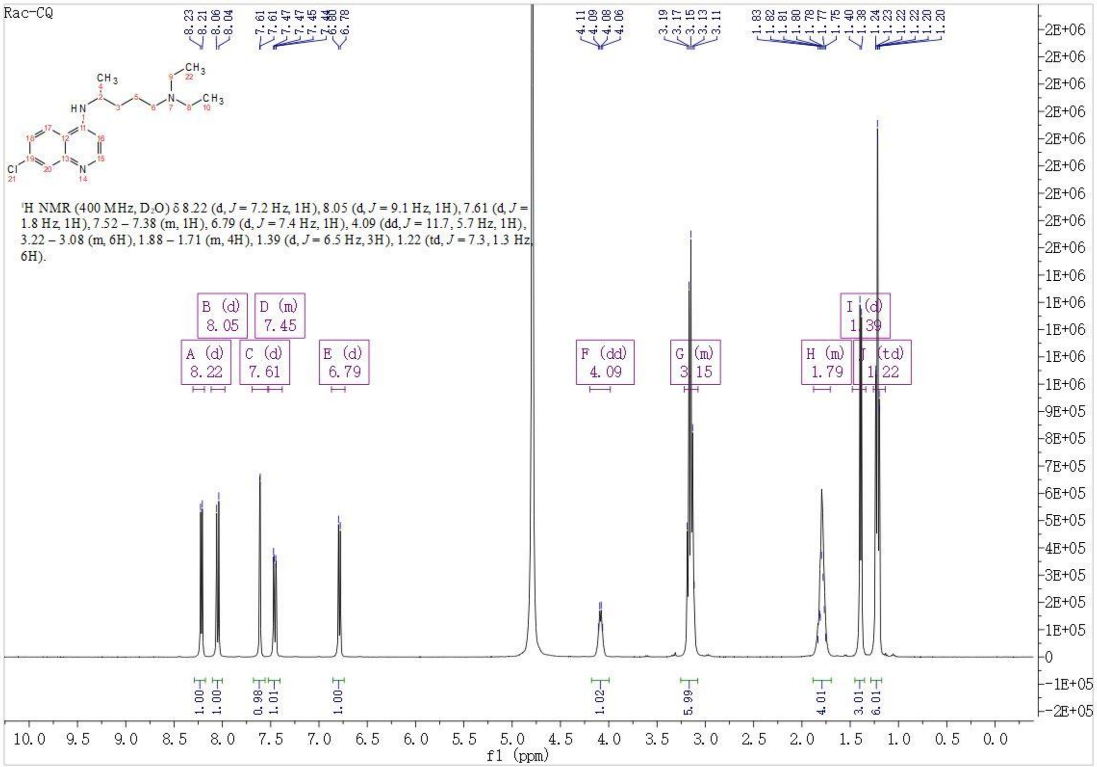


Figure S2. $^{13}C$ NMR (101 MHz, $D_2O$) of Rac-CQ diphosphate.

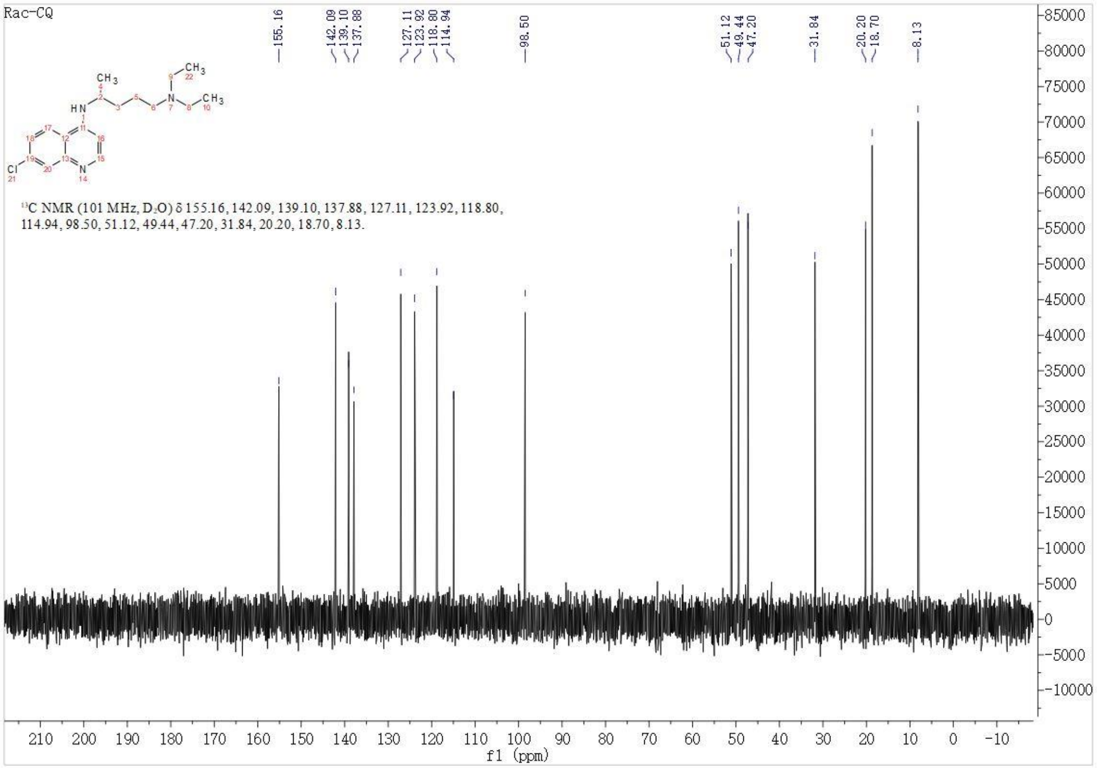

Figure S3. $^{1}H$ NMR (400 MHz, $D_2O$) of Rac-HCQ sulfate.

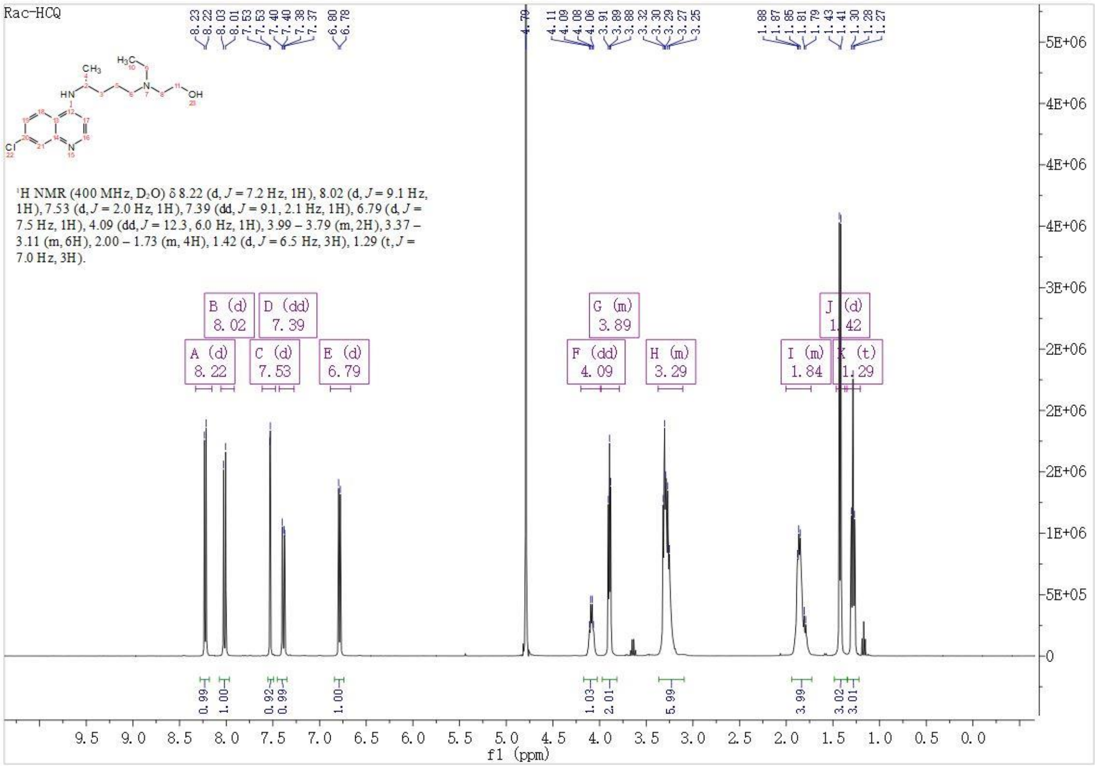

Figure S4. $^{13}C$ NMR (101 MHz, $D_2O$) of Rac-HCQ sulfate.

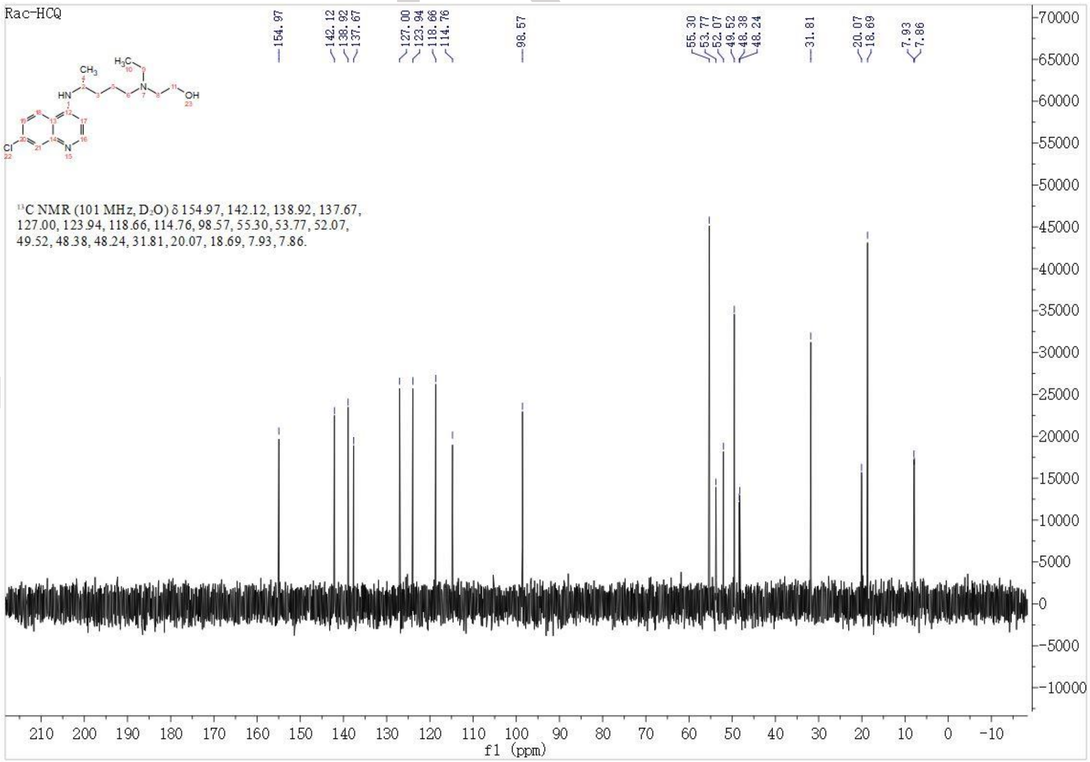

Figure S5. $^{1}H$ NMR (400 MHz, $D_2O$) of R-CQ diphosphate.

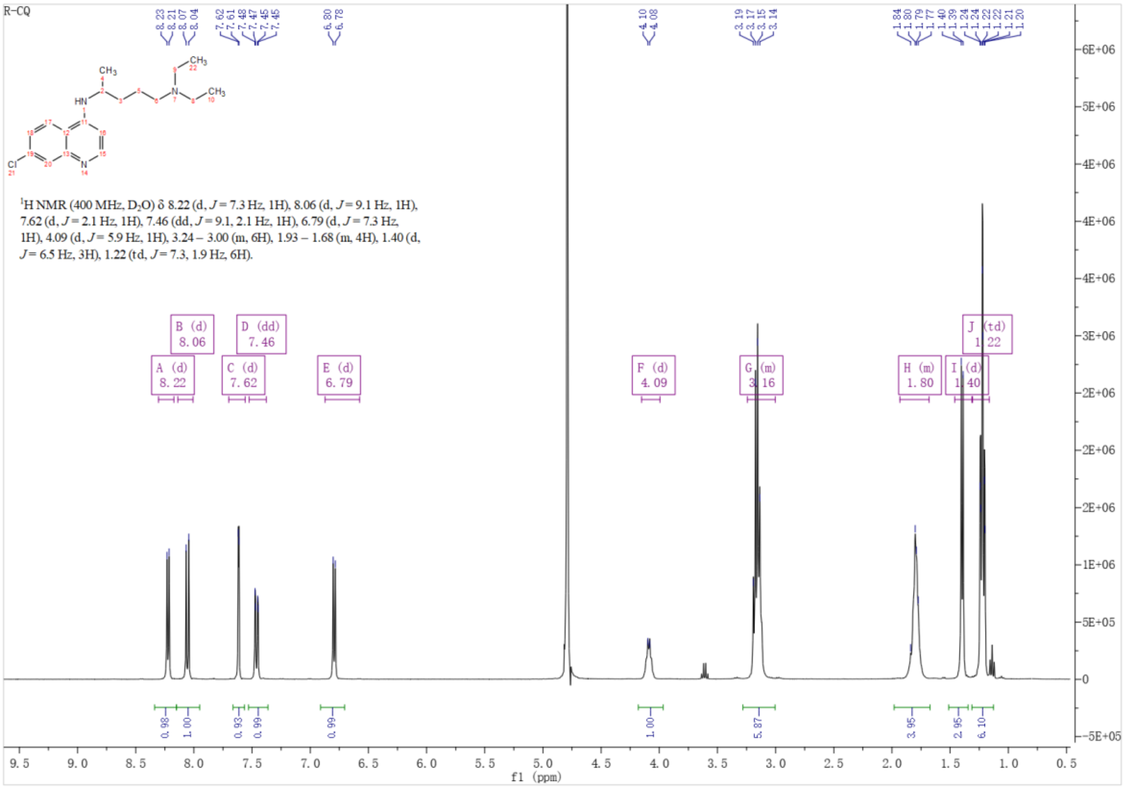


Figure S6. $^{13}C$ NMR (101 MHz, $D_2O$) of R-CQ diphosphate.

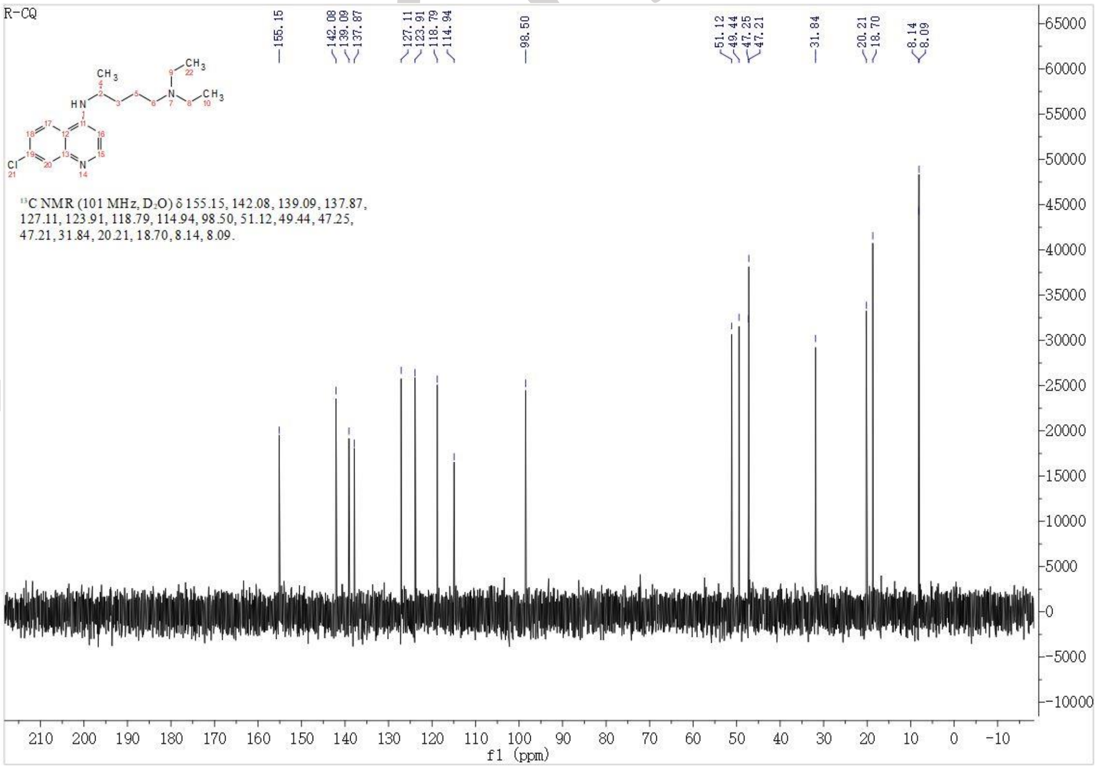

Figure S7. $^{1}H$ NMR (400 MHz, $D_2O$) of S-CQ diphosphate.

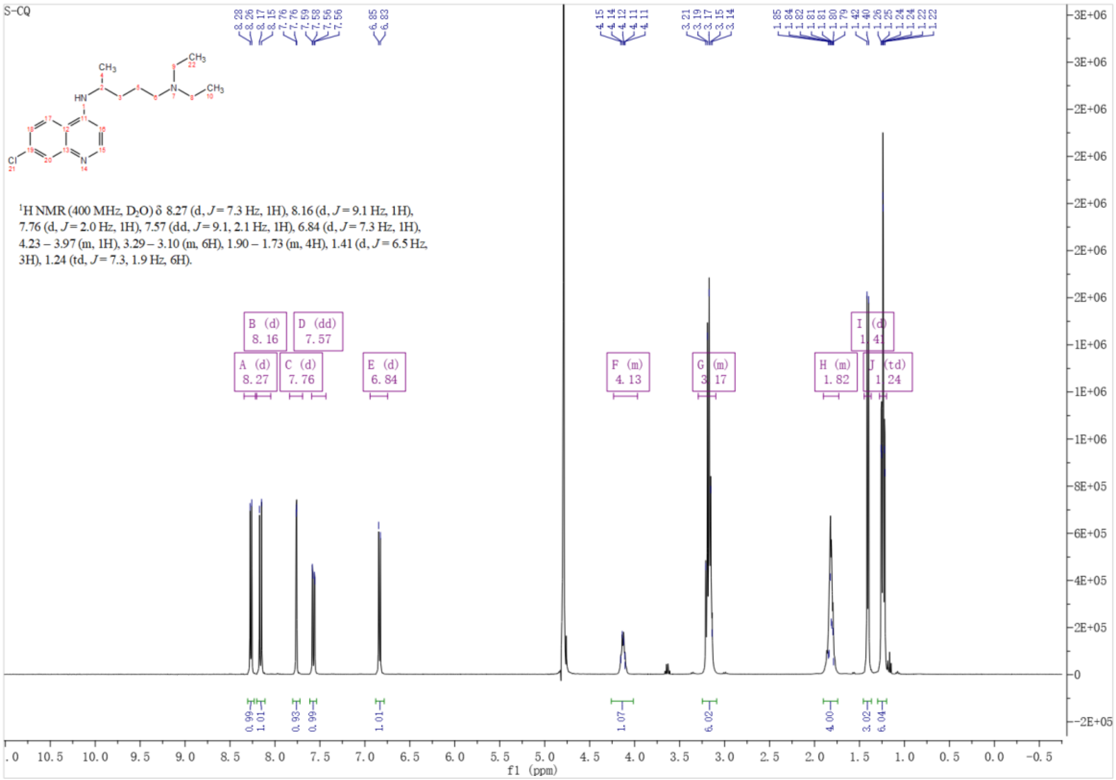


Figure S8. $^{13}C$ NMR (101 MHz, $D_2O$) of S-CQ diphosphate.

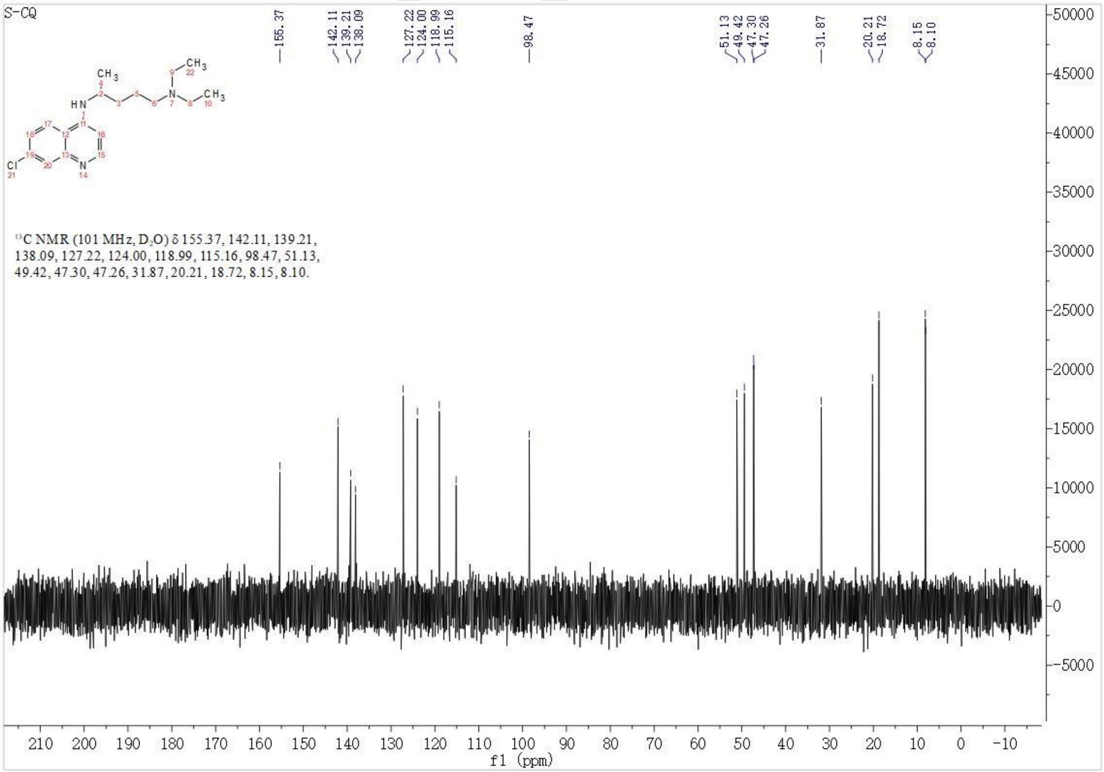

Figure S9. $^{1}$H NMR (400 MHz, $D_2O$) of R-HCQ sulfate.

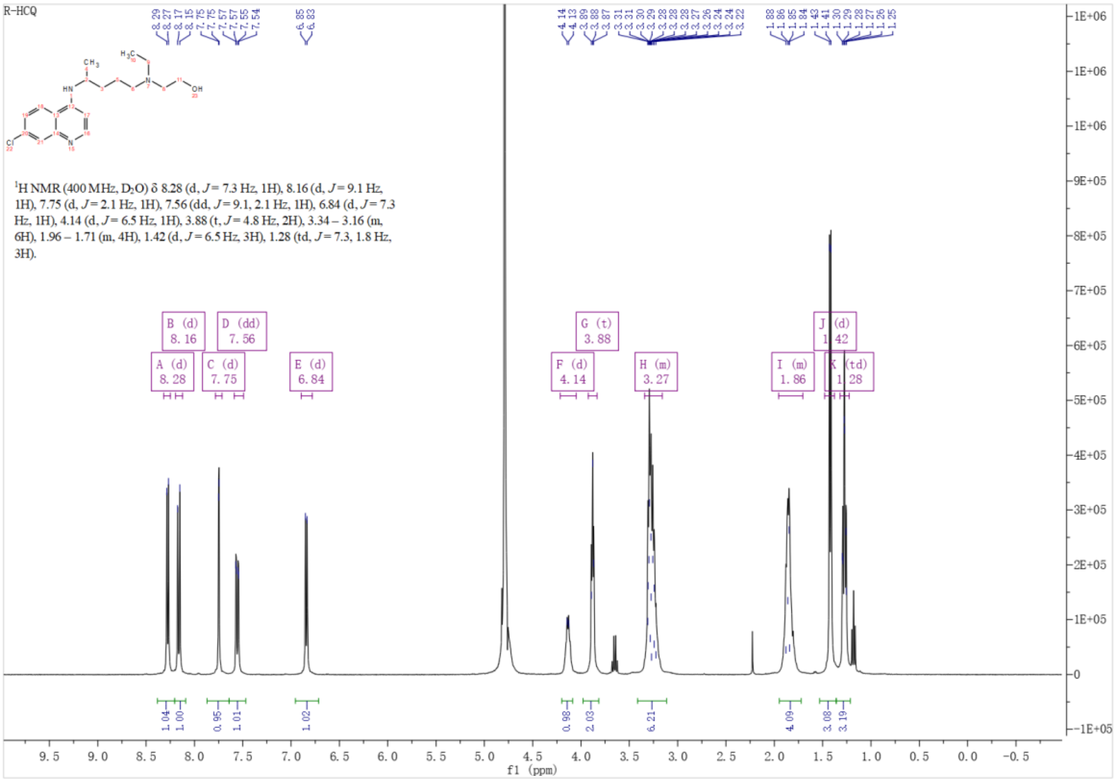


Figure S10. $^{13}$C NMR (101 MHz, $D_2O$) of R-HCQ sulfate.

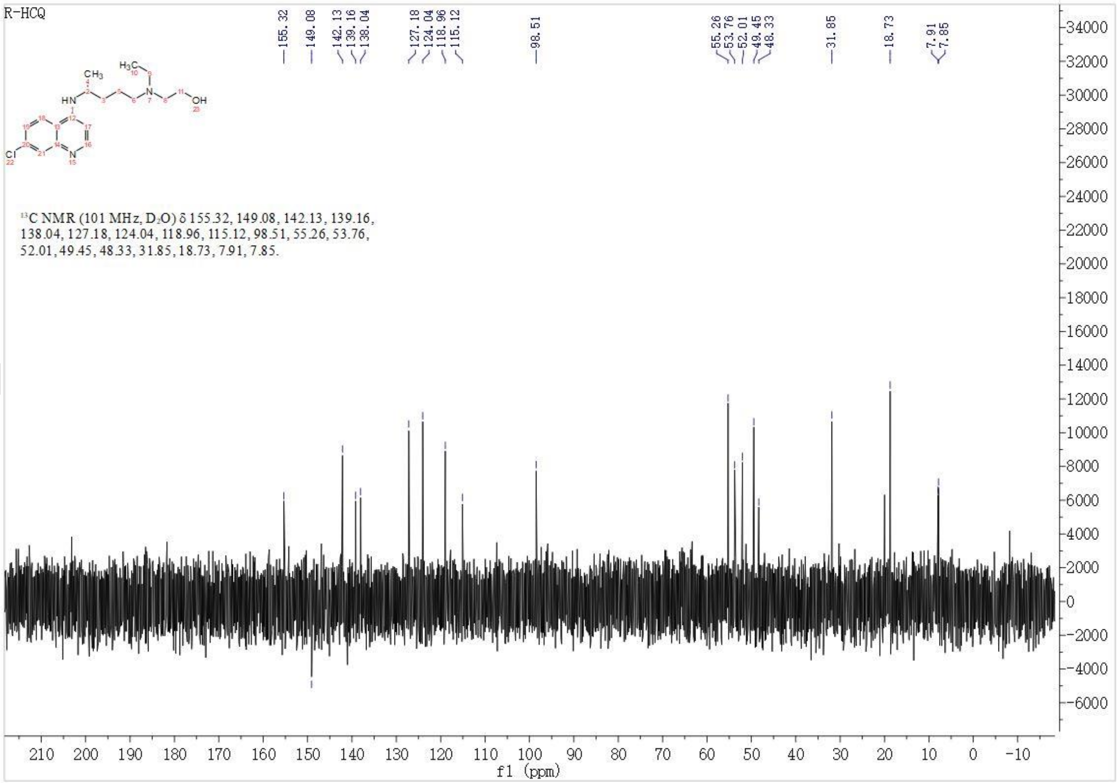

Figure S11. $^{1}$H NMR (400 MHz, $D_2O$) of S-HCQ sulfate.

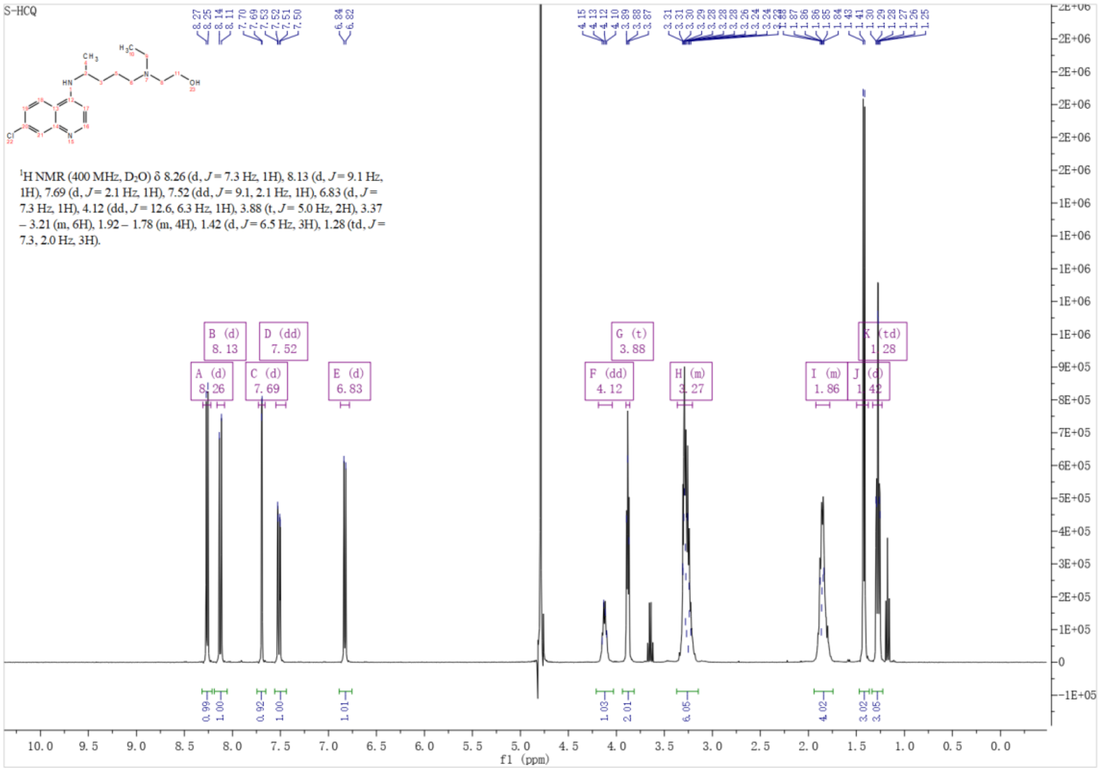


Figure S12. $^{13}$C NMR (101 MHz, $D_2O$) of S-HCQ sulfate.

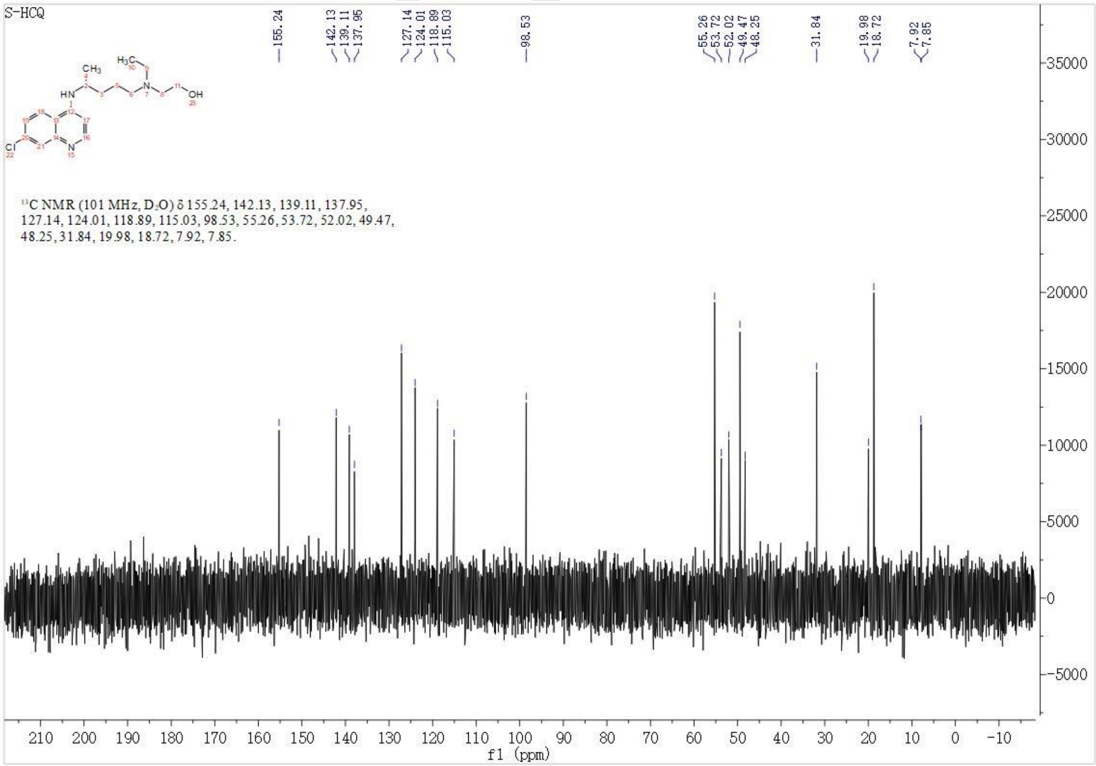

Figure S13. Antiviral activities (second batch)

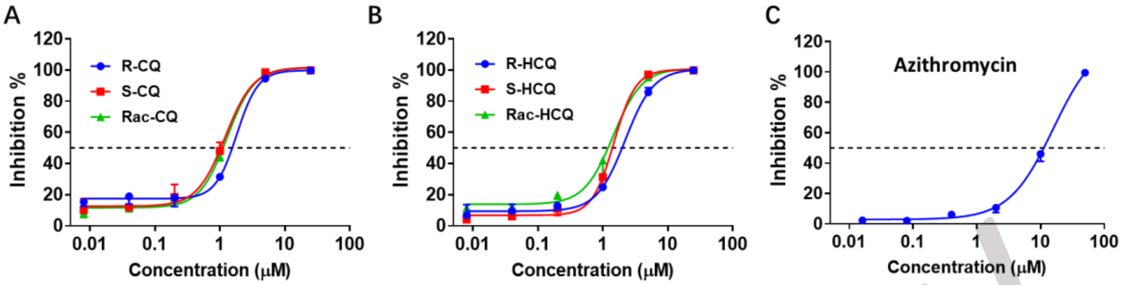


| | R-CQ | S-CQ | Rac-CQ |
|---|---|---|---|
| $IC_{50}$ (μM) | 1.818 | 1.208 | 1.286 |

| | R-HCQ | S-HCQ | Rac-HCQ |
|---|---|---|---|
| $IC_{50}$ (μM) | 2.21 | 1.483 | 1.432 |

| | Azithromycin |
|---|---|
| $IC_{50}$ (μM) | 15.75 |

The antiviral activities of racemic and enantiomeric chloroquine diphosphate (A) and hydroxychloroquine sulfate (B), as well as azithromycin against SARS-CoV-2 *in vitro*. Vero E6 cells were infected with SARS-CoV-2 (MOI = 0.05) at different concentrations: 0.008, 0.04, 0.2, 1, 5, and 25 μM for CQ and HCQ for 24 h (concentrations at 0.008, 0.04, 2, 10 and 50 μM for AZM). Data represented are the mean value of % inhibition of SARS-CoV-2 on Vero E6 cells. Experiments were performed three times for each batch, independently.

DO NOT COPY

Figure S14. Antiviral activities (statistics)

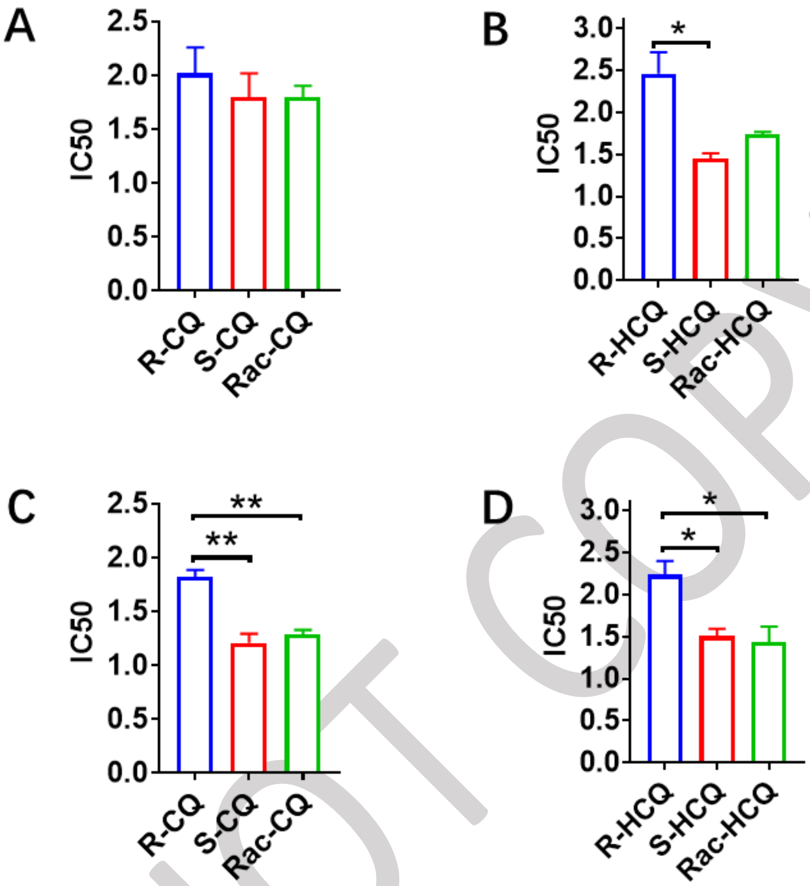


The antiviral activities of racemic and enantiomeric chloroquine diphosphate (A: batch 1, C: batch 2) and hydroxychloroquine sulfate (B: batch 1, D: batch 2) against SARS-CoV-2 *in vitro*. Vero E6 cells were infected with SARS-CoV-2 (MOI = 0.05) at different concentrations: 0.008, 0.04, 0.2, 1, 5, and 25 μM, for 24 h. Data represented are the mean value of % inhibition of SARS-CoV-2 on Vero E6 cells. Experiments were performed three times for each batch, independently. ANOVA were used to analyze differences in mean values between groups using GraphPad Prism 7. All results are expressed as mean ± standard error of the mean (SEM) and were corrected for multiple comparisons. P values of <0.05 were considered statistically significant. (*, P values of ≤0.05. **, P values of ≤0.005. ***, P values of ≤0.0005. ****, P values of ≤0.0001).

DO NOT COPY
